# Structure of PINK1 reveals autophosphorylation dimer and provides insights into binding to the TOM complex

**DOI:** 10.1101/2021.08.05.455304

**Authors:** Shafqat Rasool, Simon Veyron, Naoto Soya, Mohamed Eldeeb, Gergely L. Lukacs, Edward A. Fon, Jean-François Trempe

## Abstract

Mutations in PINK1 causes autosomal-recessive Parkinson’s disease. Mitochondrial damage results in PINK1 import arrest on the Translocase of the Outer Mitochondrial Membrane (TOM) complex, resulting in the activation of its ubiquitin kinase activity by autophosphorylation and initiation of Parkin-dependent mitochondrial clearance. Herein we report crystal structures of the entire cytosolic domain of insect PINK1. Our structures reveal a dimeric autophosphorylation complex targeting phosphorylation at the invariant Ser205 (human Ser228). The dimer interface requires insert 2, which is unique to PINK1. The structures also reveal how an N-terminal helix binds to the C-terminal extension and provide insights into stabilization of PINK1 on the core TOM complex.

## Introduction

Mutations in PINK1 (PARK6) and Parkin (PARK2) cause early-onset autosomal recessive Parkinson disease (PD) (Kitada et al., 1998; Valente et al., 2004). These two proteins function together in a mitochondrial quality control pathway (Bayne and Trempe, 2019). Upon mitochondrial damage, the serine-threonine kinase PINK1 accumulates on the surface of the mitochondria (Geisler et al., 2010; Narendra et al., 2010; Vives-Bauza et al., 2010). Following activation, PINK1 phosphorylates mitochondrially-tethered ubiquitin (Ub) at a conserved Ser65 signaling the recruitment of the RBR E3-ligase Parkin to damaged mitochondria (Kondapalli et al., 2012). Binding to phosphorylated Ub (pUb) induces the dissociation of the buried ubiquitin-like domain (Ubl) of Parkin, which can then be phosphorylated by PINK1, resulting in the activation of its ligase activity (Kazlauskaite et al., 2015; Kumar et al., 2015; Sauvé et al., 2015; Wauer et al., 2015). Activated parkin ubiquitinates OMM proteins such as Mitofusin-1 and -2 as well as VDAC, in immortalized cancer cells and neurons (Ordureau et al., 2020; Sarraf et al., 2013; Tanaka et al., 2010). Upon build-up of a critical mass of Ub chains by feed forward cycles of Ub phosphorylation, Parkin recruitment and activation, autophagy receptors such as NDP52 and OTPN (activated by TBK1 kinase) converge on the mitochondria and signal the recruitment of the autophagy machinery to engulf the damaged organelle (Heo et al., 2015; Lazarou et al., 2015).

The accumulation of PINK1 on mitochondria is regulated by a complex array of interactions and proteolytic steps. PINK1 is imported through its N-terminal mitochondrial targeting sequence (MTS) and successively cleaved by the mitochondrial processing peptidase (MPP) and the Presenilin-associated rhomboid-like (PARL) and AFG3L2 proteases (Greene et al., 2012; Jin et al., 2010). This creates a processed 52 kDa species that becomes a substrate for the N-end rule E3 ligases Ubr1, Ub2 and Ubr4 in the cytosol (Yamano and Youle, 2013). PINK1 thus undergoes rapid turnover by the proteasome via ubiquitination (Fedorowicz et al., 2014), which results in low steady-state levels. However, when mitochondria are exposed to stress such as depolarization or unfolded proteins, PINK1 is no longer cleaved and accumulates as a full-length 64 kDa protein bound to the Translocase of the Outer Mitochondrial Membrane (TOM) complex (Lazarou et al., 2012). This coincides with its activation through autophosphorylation, followed by Ub phosphorylation (Koyano et al., 2013; Okatsu et al., 2012).

In the last three years, the structure of PINK1’s kinase domain and the mechanism of Ub and Parkin phosphorylation have been elucidated using insect orthologs of PINK1 from *Tribolium castaneum* (beetle) and *Pediculus humanus corporis* (lice) (Gladkova et al., 2017; Kumar et al., 2017; Rasool et al., 2018; Schubert et al., 2017). PINK1 orthologs have a conserved bilobular kinase domain with canonical features found in other protein kinases. It also houses three unique inserts in the N-lobe as well as a C-terminal helical bundle that stacks on top of the C-lobe, known as the C-terminal extension (CTE). Crystal structures and NMR studies revealed how PINK1 bind its substrates and provided insights into the conformational activation of PINK1(Rasool and Trempe, 2018). Importantly, insert 3 and the P-loop in the N-lobe form the key docking sites for the Ile44 binding patch of ubiquitin, while a conserved stretch of residues in the activation loop (A-loop, a.a. 402-414 in human PINK1) interact with the target Ser65 loop (Schubert et al., 2017). Slippage of the β5 strand has been proposed to allow the access of the Ser65 by the PINK1 active site (Gladkova et al., 2017).

The N-lobe of the kinase domain also harbors a unique autophosphorylation site (Ser228 in hPINK1; Ser205 in TcPINK1) located in the ‘C-loop’ connecting β3 and the αC helix of the N-lobe (Rasool and Trempe, 2018). We have shown that phosphorylation at this site occurs in *trans*, a step that is critical for Ub binding to PINK1 (Rasool et al., 2018). Comparison of the apo and ubiquitin-bound phosphorylated PINK1 structure as well as data from hydrogen-deuterium exchange mass spectrometry (HDX-MS) suggests that phosphorylation of Ser228 enables a remodeling of the N-lobe of the kinase domain to bring about insert 3 refolding and accommodating Ub at the active site (Rasool and Trempe, 2018; Schubert et al., 2017). Intriguingly, mutants of the PINK1 autophosphorylation site retain kinase activity, showing that phosphorylation at this site primes PINK1 specifically for Ub binding (Rasool et al., 2018). In line with this, insert 3 deletion abrogates Ub phosphorylation but does not affect *trans* autophosphorylation (Kumar et al., 2017). Since *trans* autophosphorylation is the prerequisite for Ub and Ubl phosphorylation, understanding its molecular mechanism is key to fully uncovering the function of PINK1.

In recent years the mechanism of PINK1 activation on damaged mitochondria has been under investigation. PINK1 localizes on damaged mitochondria in complex with components of the TOM complex (TOM40, TOM20, TOM22 and TOM70), forming a ∼700-800 kDa complex detected by native gels (Lazarou et al., 2012; Okatsu et al., 2013). PINK1 probably exists as a dimer in this complex, which coincides with activation via autophosphorylation (Okatsu et al., 2013). Later studies identified TOM7 as a key component allowing for the stabilization of PINK1 on the mitochondria (Hasson et al., 2013; Sekine et al., 2019). The stabilization of PINK1 also depends on a stretch of negatively charged residues at the beginning of a putative N-terminal (NT) helix, a ∼30 a.a. linker located between the TM segment and the kinase domain that harbors PD mutations (Sekine et al., 2019). Despite these observations, the localization of PINK1 is far from being fully understood as neither the structure of the NT linker nor the precise interactions made by the above mentioned negatively charged residues are known. The precise arrangement of activated PINK1 in the TOM complex and whether this step is anchored to autophosphorylation also remain to be known.

Herein we present our findings about the autophosphorylation mechanism of PINK1 and the role of the NT linker in mediating key inter and intra-molecular interactions that allow for its localization on the TOM complex. We have solved the crystal structure of the entire C-terminal domain of PINK1 from *Tribolium castaneum* (TcPINK1) in both dephosphorylated and phosphorylated forms, which reveal a symmetric *trans* autophosphorylation complex with the acceptor Ser205 in position to receive a phosphate from ATP. The interface is conserved and requires insert 2. Our structures also show that the NT linker is composed of a single long α-helix that interacts with the αK helix in the CTE. Finally, molecular modeling using our crystal structures and recently published cryoelectron microscopy (cryoEM) structures of the TOM complex predicts interactions between the NT helix and TOM7.

## Results

### Crystal structures of PINK1 reveal a dimeric *trans* autophosphorylation complex

Wild-type (WT) TcPINK1 phosphorylates kinase-dead (D337N) TcPINK1 efficiently, with a catalytic efficiency k_cat_/K_m_ of 6.5 μM^-1^ min^-1^ (Figure 1A), compared with 0.22 μM^-1^ min^-1^ and 0.05 μM^-1^ min^-1^ for Parkin Ubl and Ub, respectively (Rasool et al., 2018). TcPINK1 must therefore form a complex in order to phosphorylate itself in *trans*. To capture this complex, we sought to produce fully non-phosphorylated active PINK1. Overexpression in *E. coli* leads to massive spurious autophosphorylation (Rasool et al., 2018), and thus we co-expressed WT TcPINK1 with the lambda phage phosphatase to produce completely dephosphorylated protein after purification (Figure 1B). Incubation of the protein with Mg^2+^-ATP *in vitro* led to the formation of homogenously monophosphorylated TcPINK1, with a single phosphorylation at Ser205 (Figure 1B and Suppl. Table S1). However, the resulting protein had low solubility and thus we introduced six solubilizing mutations in solvent-exposed aromatics. Our crystallization construct spans a.a. 121-570 (TcPINK1^121-570^-6arom), which includes the NT linker. The non-phosphorylated and phosphorylated forms of this construct crystallized in the same space group in the presence of AMP-PNP and diffracted X-rays to 3.0 Å and 2.8 Å resolution, respectively (Table 1). The two structures are highly similar (RMSD 0.4 Å), with an asymmetric unit containing a single protein chain (Figure 1C and Suppl. Figure S1A). The region spanning a.a. 150-570 superposes well on the previous TcPINK1 structures (RMSD ∼0.6 Å with PDB 5OAT and 5YJ9) showing that the overall structure is similar. However, crystallographic symmetry analysis in both our structures reveals a novel face-to-face dimer, forming an interface with a buried surface area of 4800 Å^2^ (Figure 1D). The structure of the dimer is unique compared to other dimeric kinase autophosphorylation complexes (Suppl. Figure S1B). In this dimeric assembly, the exposed acceptor Ser205 (or phospho-Ser205) from one subunit interacts with the catalytic loop Asp337 from another subunit in a reciprocal manner. This strongly suggests that this dimer corresponds to the *trans* autophosphorylation complex.

**Table 1.** X-ray crystallography data collection and refinement statistics.

|  | **TcPINK1<sup>121-570</sup>** | **Phospho-TcPINK1<sup>121-570</sup> + AMP-PN** |
| --- | --- | --- |
| **Wavelength (Å)** | 0.97949 | 0.97949 |
| **Resolution range (Å)** | 49.58-3.00 (3.18-3.00) | 45.67-2.80 (2.92-2.80) |
| **Space group** | P6<sub>1</sub>22 | P6<sub>1</sub>22 |
| **Unit cell dimensions (Å)** | 52.9137 52.9137 545.362 | 52.736 52.736 547.944 |
| **Unit cell angles (deg)** | 90 90 120 | 90 90 120 |
| **Total reflections** | 169985 (19995) | 334179 (16648) |
| **Unique reflections** | 10223 (1589) | 9568 (478) |
| **Multiplicity** | 16.4 (12.6) | 34.9 (34.8) |
| **Completeness (%)** | 100 (100) | Spherical 76.7 (33.0)<br>Ellipsoidal 93.4 (100) |
| **Mean I/sigma(I)** | 6.1 (1.1) | 10.6 (0.7) |
| **Wilson B-factor** | 62.83 | 34.9 |
| **R-merge** | 0.737 (3.613) | 0.533 (2.726) |
| **CC<sub>1/2</sub>** | 0.989 (0.627) | 0.996 (0.841) |
| **Reflections used for R-free** | 478 | 445 |
| **R-work** | 0.2538 | 0.2469 |
| **R-free** | 0.3062 | 0.3065 |
| **# non-hydrogen atoms** | 3058 | 3335 |
| **macromolecules** | 3014 | 3188 |
| **ligands** | 10 | 87 |
| **water** | 34 | 60 |
| **Protein residues** | 399 | 414 |
| **RMS (bonds, Å)** | 0.003 | 0.003 |
| **RMS (angles, deg)** | 0.587 | 0.548 |
| **Ramachandran favored (%)** | 96.66 | 94 |
| **Ramachandran allowed (%)** | 3.08 | 6 |
| **Ramachandran outliers** | 0.26 | 0 |
| **Clashscore** | 3.71 | 5 |
| **Average B-factor** | 72.81 | 28.38 |
| **macromolecules** | 72.85 | 27.94 |
| **ligands** | 99.78 | 47.73 |
| **solvent** | 61.02 | 23.52 |

Statistics for the highest-resolution shell are shown in parentheses.

**Figure 1.**
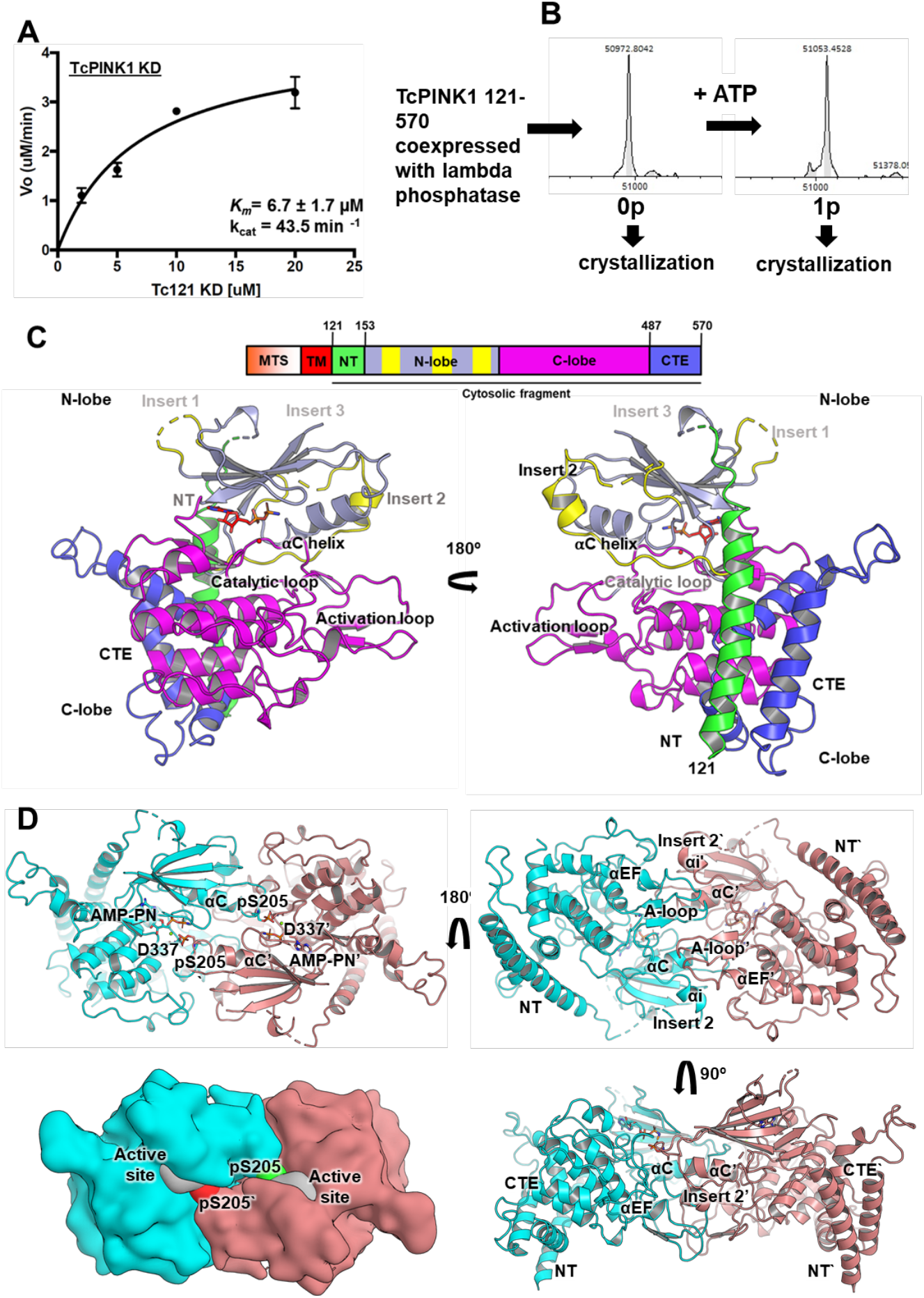
Crystal structures of PINK1 reveal a dimeric *trans* autophosphorylation complex. **(A)** Enzyme kinetics of the phosphorylation of kinase dead (KD) TcPINK1^121-570^ D337N by WT GST-TcPINK1^121-570^. Phosphorylation assays were performed for 1 min using different concentrations of KD and 0.1 µM WT enzyme and were analyzed using intact mass spectrometry followed by Michaelis-Menten modelling. The given graph represents global fit to data collected from 2 sets of reactions performed independently. Bars represent mean ± SD (n=2). **(B)** Intact mass spectra of TcPINK1^121-570^ crystallization construct co-expressed with lambda phosphatase before (left) and after phosphorylation with ATP. **(C)** Structure of the asymmetric unit for Ser205-phosphorylated TcPINK1^121-570^. **(D)** Crystallographic symmetry reveals a two-fold symmetric face-to-face dimer that captures the autophosphorylation complex with the phospho-Ser205 from each monomer (cyan and pink) interacting with the catalytic Asp337 from the other chain (top left). Top right and bottom right panels show rotated poses, while the bottom left panel shows a surface representation of the top left pose.

Close inspection of the dimer interface highlights interactions mediated by the C-loop and the αi helix located in insert 2 (Figure 2A). The conserved Lys339 in the catalytic loop stabilizes this interaction. Additionally, and similarly to the loop harboring Ser65 in Ub (Schubert et al., 2017), the C-loop harboring Ser205 in the acceptor subunit interacts with the backbone of a.a. 383-385 in the A-loop of the donor (Figure 2B). Additional stabilization is provided by hydrophobic contacts between the conserved Tyr429 in the C-lobe and Ile203 in the C-loop of the opposing subunit (Figure 2B). Consistent with these observations, TcPINK1 mutants K339A or Y429A exhibited a reduced ability to autophosphorylate *in vitro* at Ser205, as observed using intact mass spectrometry (Figure 2C,D and Suppl. Figure S2A). Additionally, we noted a hydrogen bond between Thr386 and Ser207. While Ser207 (Ser204 in PhPINK1) phosphorylation was suggested to be important for Ub binding (Schubert et al., 2017), TcPINK1 S207A was able to phosphorylate Ub, suggesting Ser207 is dispensable (Kumar et al., 2017). Based on our structure, the introduction of a bulkier phosphate or carboxylic acid group at Ser207 creates a clash with the side chain of Thr386 in the opposite subunit, and thus would disrupt the interface. Indeed, we find that mutation of TcPINK1 Ser207 to a phosphomimetic aspartate results in a complete loss of autophosphorylation activity at Ser205 (Figure 2C,D, Suppl. Figure S2A). Overall, this data is consistent with the observed dimer being the genuine autophosphorylation complex.

**Figure 2.**
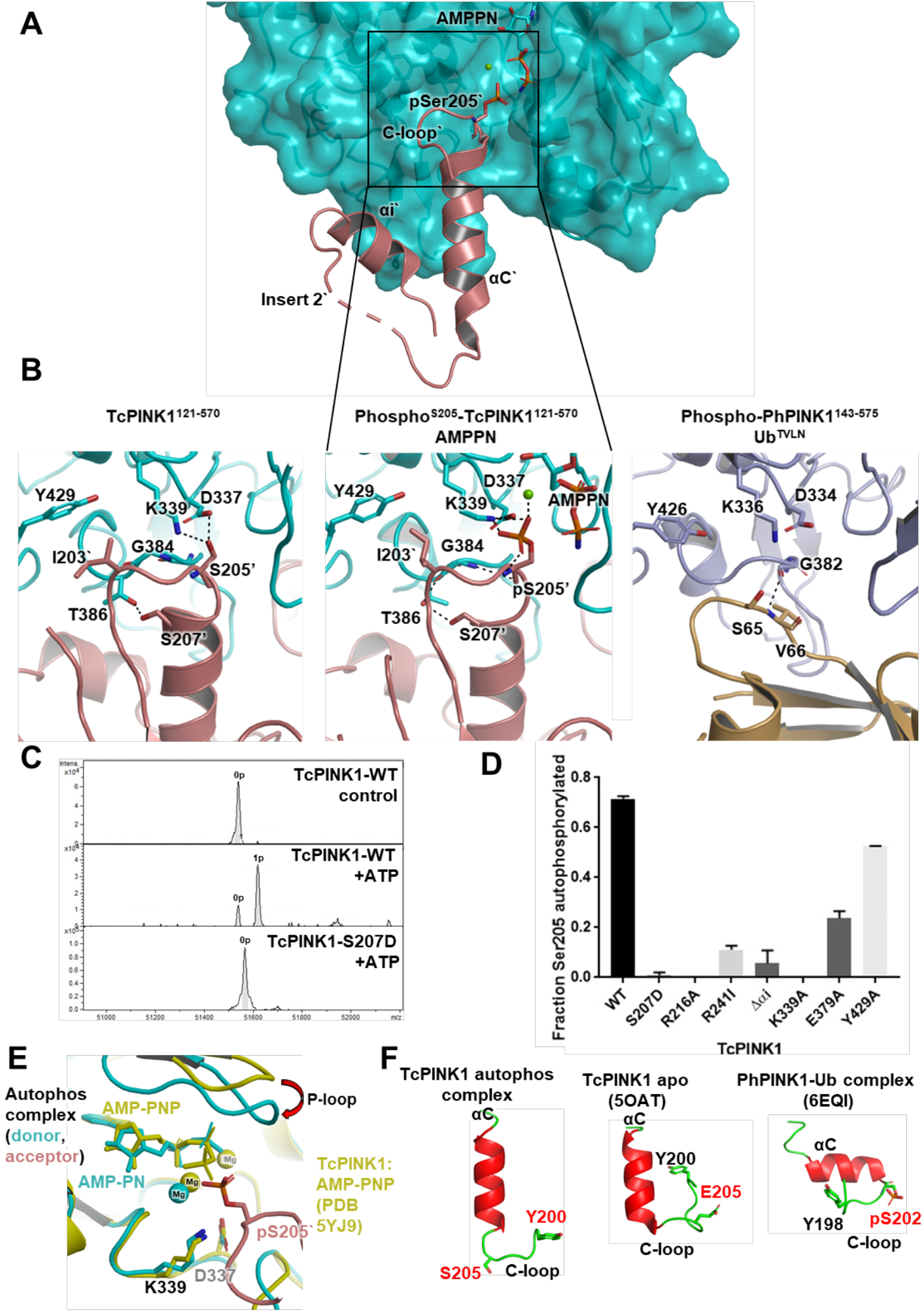
Interactions at the active site in the autophosphorylation complex. **(A)** The αC helix and the αi helix (from insert 2) from one monomer (pink) in the complex interact with the other monomer (cyan; surface). **(B)** Interactions at the active site in the non-phosphorylated (left) and phosphorylated (middle) autophosphorylation complex structures, and previously published PhPINK-Ub^TVLN^ complex structure (PDB: 6EQI). **(C)** Intact mass spectra of the phosphorylation reactions performed by incubating 1 µM dephosphorylated WT or S207D TcPINK1^121-570^ with ATP for 30 seconds at 30°C. **(D)** Fractional levels of phosphorylation achieved during phosphorylation assays by WT of autophosphorylation interface mutants. Bars represent mean ± SD (n=2). **(E)** Comparison (overlay) of the P-loop and catalytic site conformation of the previously published AMP-PNP bound TcPINK1 structure (yellow) to phospho-Ser205 autophosphorylation complex. **(F)** Side-by-side comparison between the conformations of αC helix and the C-loop (containing Ser205) between the autophosphorylation complex (this study), TcPINK1 apo structure (5OAT) and the PhPINK-Ub^TVLN^ complex.

Key differences are noted in the conformation of canonical kinase structural elements upon comparison with other PINK1 structures. Our phosphorylated TcPINK1 structure showed clear electron density for a nucleotide in the active site (Suppl. Figure S2B). The P-loop (a.a. 170-175), which is involved in nucleotide binding, moves inwards and closer to the active site in our structure compared to the previously published nucleotide-bound structure (Okatsu et al., 2018), reminiscent of a protein conformation primed for catalyzing phosphoryl transfer (Figure 2E). The αC helix, which is kinked in the Ub-bound PhPINK1 structure, adopts a straight conformation similar to the apo S205E structure (Figure 2F). However, the C-loop that precedes the αC helix is displaced outwards relative to all the other TcPINK1 structures, thus exposing Ser205 to the surface. This unique conformation is critical to allow Ser205 in the acceptor subunit to reach the active site in the donor subunit, thus allowing *trans* phosphorylation to take place.

### Insert 2 is required for autophosphorylation

Multiple interactions between the two protein molecules stabilize the autophosphorylation complex, distal to the Ser205 phosphorylation site (Figure 3A). These interactions involve the αi helix (a.a. 233-241) in insert 2, the activation loop (A-loop) and the αEF helix. The A-loop segment 375-381 protrudes into the cavity between the αC helix, insert 2 and the A-loop of the opposing subunit (Figure 3B). Glu379 (A-loop) lies at the apex of this protrusion and forms salt bridges with Arg241 in insert 2 and Arg216 in the αC helix (Figure 3B). The αi helix stacks against residues 395-400, just downstream of the APE motif that marks the end of the activation loop. Trp233 in αi forms hydrophobic contacts with Cys395 and residues Phe437-Asn438. In the apo and Ub-bound PINK1 structures, the αi helix forms intramolecular interactions (this helix is not visible in the previous published AMP-PNP bound structure). A superposition of the αi helices of the three structures shows that relative to the apo and Ub-bound structures, the αi helix in our structure is fully displaced outwards, exposing Trp233 which then interacts with the opposing subunit (Figure 3C). Consistent with these observations, R216A, R241I, E379A or Δ231-242 (Δαi) mutants of TcPINK1 strongly reduced the ability of TcPINK1 to autophosphorylate in vitro (Figure 2D and Suppl. Figure S2). The importance of these distal interactions in the formation of this complex and *trans* autophosphorylation is highlighted by our observation that TcPINK1 does not phosphorylate a synthetic peptide spanning TcPINK1 a.a. 199-209, which contains Ser205 and residues proximal to the active site in the acceptor subunit (Suppl. Figure S3). Distal interactions are therefore critical for autophosphorylation.

**Figure 3.**
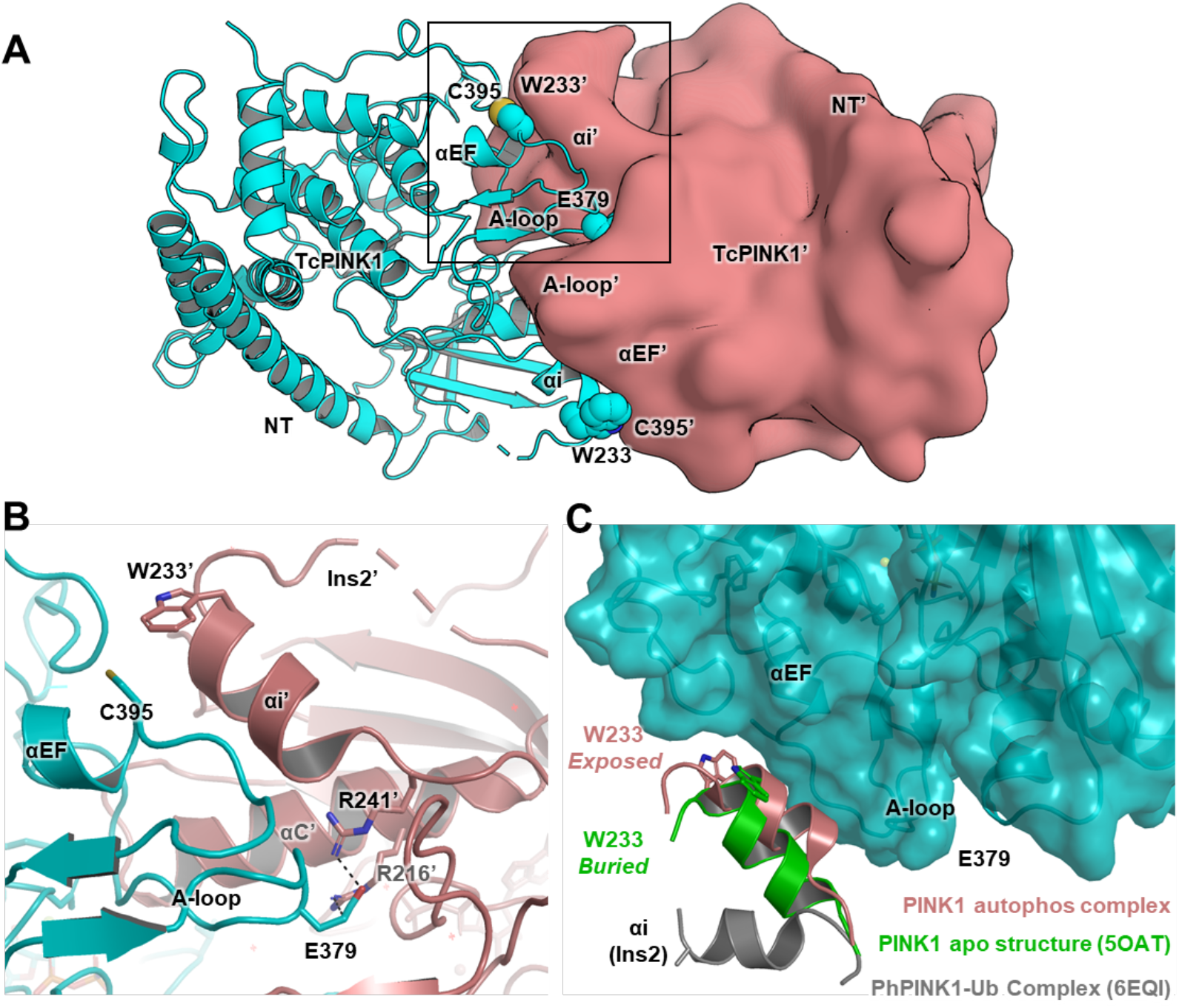
Insert 2 is an essential part of the autophosphorylation interface. **(A)** 180° rotated pose of autophosphorylation complex (as in Figure 1D; top left) depicting the two monomers (cyan cartoon, pink surface) and elements of the kinase mediating distal interactions. **(B)** Zoomed version of the inset in A showing the interactions between Glu379 in the A-loop with Arg241 in αi and Arg216 in the αC helix of the opposing monomer. **(C)** Comparison of the conformation of αi helix between autophosphorylation complex (this study), the TcPINK1 apo structure and the PhPINK1:Ub^TVLN^ complex. In the autophosphorylation complex, Trp233 in the αi flips outwards to become exposed, while it is buried in the TcPINK1 apo (PDB 5OAT).

### The autophosphorylation interface is distinct from the ubiquitin binding site

Comparison of the autophosphorylation dimer with the PhPINK1:Ub^TVLN^ complex (Schubert et al., 2017) shows that the two interfaces overlap, implying that the two substrates are mutually exclusive, but yet involve different elements (Figure 4A). In particular, interactions mediated by Arg216, Arg241 (αi helix), Glu379 and the αEF helix in the autophosphorylation complex are not involved in binding Ub. To validate this observation, we tested the ability of our TcPINK1 mutants to phosphorylate the Parkin Ubl domain. Since autophosphorylation at Ser205 is an essential prerequisite for Ub/Ubl binding, all mutants were autophosphorylated for an extensive period of time until all mutants were fully phosphorylated, prior to conducting Ubl phosphorylation reactions (Suppl. Figure S4). Phospho-K339A was mildly compromised in Ubl phosphorylation (Figure 4B), which is consistent with Lys339 playing a role in catalyzing phosphate transfer. Likewise, the phosphorylated Y429A mutant was also impaired in Ubl phosphorylation. The equivalent residue in PhPINK1 (Tyr427) is in close proximity to Ub-Glu64 in the PhPINK1:Ub^TVLN^ complex and thus is likely involved in both phosphorylation steps. On the other hand, phospho-TcPINK1 distal mutants R216A, R241I, E379A or Δ231-242 (Δαi) all phosphorylated the Ubl to the same extent as phospho-WT, whereas the same mutants have impaired autophosphorylation activity (Figure 2D). These results highlight the distinct nature of the autophosphorylation interface.

**Figure 4.**
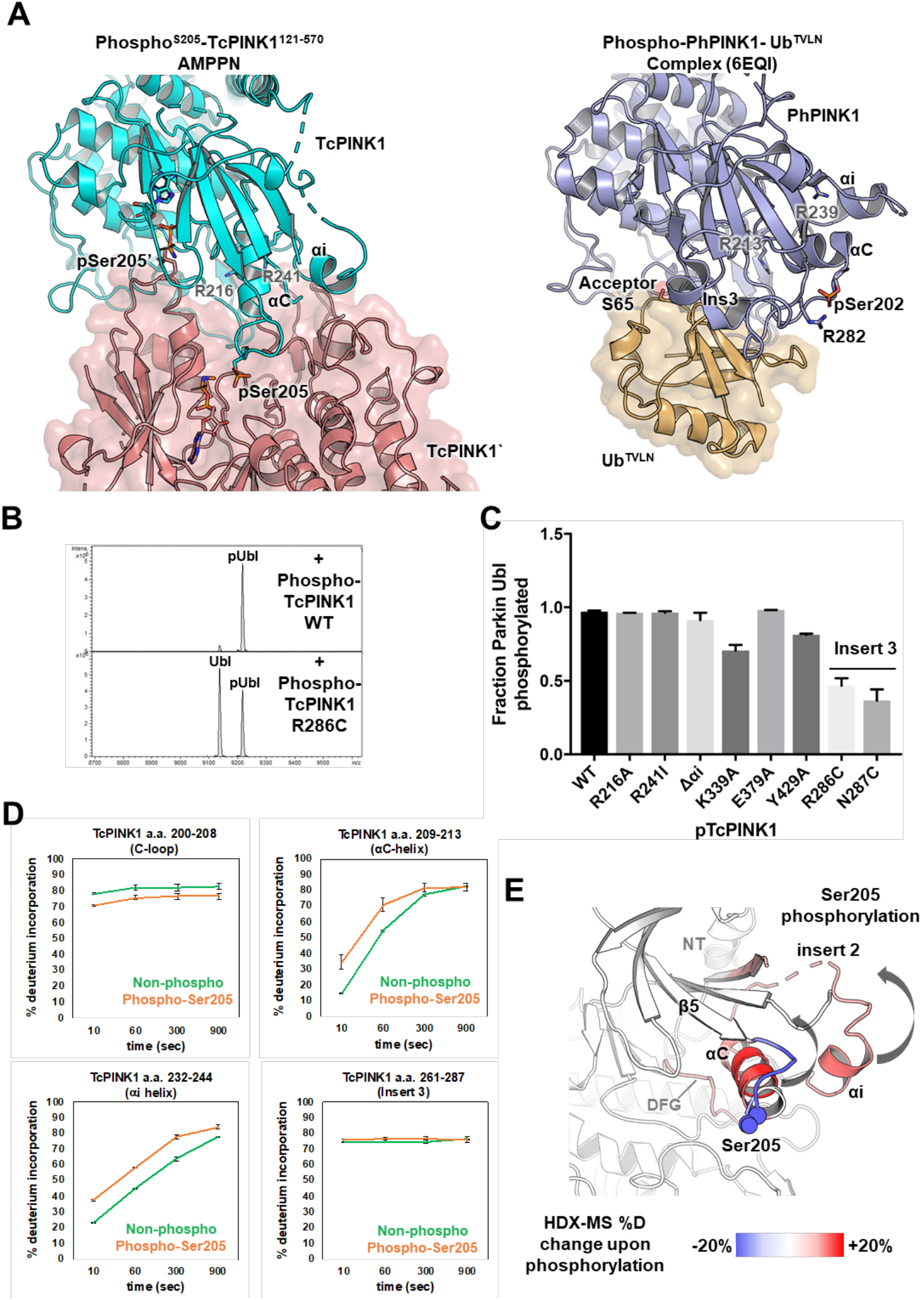
The Ub-binding and autophosphorylation interfaces are distinct. **(A)** Side-by-side comparison of the TcPINK1 autophosphorylation complex (left) and the PhPINK1-Ub^TVLN^complex (right), showing the differences in the conformation and engagement of the αC and αi helices. Insert 3 is folded and present at the ubiquitin interface in PhPINK1-Ub^TVLN^, but is disordered in the autophos complex. **(B)** Intact mass spectra of phosphorylation assays of 60 µM rat Parkin Ubl (1-76) with Ser205-phosphorylated WT or R286C TcPINK1^121-570^. **(C)** Fraction of total rat Parkin Ubl phosphorylated by Ser205-phosphorylated WT or mutant TcPINK1^121-570^. Bars represent mean ± SD (n=2). **(D)** % deuterium uptake of peptides from different regions of the kinase for non-phosphorylated and Ser205-phosphorylated WT TcPINK1^121-570^. Bars represent mean ± SD (n=3). **(E)** Difference in % deuterium uptake (between non-phosphorylated and Ser205-phosphorylated TcPINK1^121-570^) mapped onto the structure of TcPINK1. Arrows represent the conformational change that the αC and αi helices would undergo to go towards a Ub-binding competent state.

In the Ub-bound state (Schubert et al. 2017), the N-terminal region of insert 3 in PhPINK1 interacts with Ub while Arg282 and Asn283 (Arg286 and Asn287 in TcPINK1) at the C-terminus stabilize the phosphate on Ser202 (Ser205 in Tc). Insert 3 deletion in TcPINK1 (Δ261-270) is defective in Ub phosphorylation, but can still autophosphorylate (Kumar et al., 2017). In both our autophosphorylation complex structures, insert 3 (a.a. 260-282) is disordered. Arg286 and Asn287 are visible near the Ser205 loop, but do not contribute to the dimeric interface. Indeed, the R286C and N287C mutants exhibited no change in autophosphorylation activity compared to WT (Suppl. Figure S2). However, consistent with the role of insert 3 in binding Ub, phosphorylated R286C or N287C mutants were impaired in Ubl phosphorylation compared with phospho-WT (Figure 4B). This confirms that insert 3 is dispensable for PINK1 autophosphorylation but required for Ub/Ubl phosphorylation.

Furthermore, while the αC and αi helices are critical for autophosphorylation complex formation, we also observed that these two elements undergo a major concerted conformational change to bind Ub (Figure 4A). In a fully extended conformation, the αC helix as observed in our autophosphorylation complex or apo structure would clash with Ub (Rasool and Trempe, 2018). The αC helix thus adopts a kinked conformation to accommodate Ub, but also to move pSer202/205 towards insert 3 to fold this element for Ub binding. In doing so, the αi helix moves inward as it maintains mutual contacts with the kinked αC helix. In support of this model, hydrogen-deuterium exchange mass spectrometry (HDX-MS) show an increased deuterium uptake in the αC and αi helices upon Ser205 phosphorylation, and a decreased uptake in the C-loop (Figure 4 D,E and Suppl. Figure S5). However, no change is observed in peptides from insert 3, suggesting that folding may require engagement of Ub or Ubl substrate. Thus, phosphorylation at Ser205 induces specific conformational changes in the αC and αi helices, which prime PINK1 for Ub/Ubl binding.

### The autophosphorylation interface is required for activation of human PINK1

Autophosphorylation at Ser228 has been shown to be important for the activation of human PINK1 in cells (Okatsu et al., 2012; Rasool et al., 2018). Sequence analysis shows that Ser228 is invariant across PINK1 orthologs (Figure 5A and Suppl. Figure S6). Thus we hypothesized that human and insect PINK1 also autophosphorylate through a conserved interface. To test this, we transfected PINK1 knock-out (KO) U2OS cells with WT or mutants PINK1 expressed from a weakened CMV promoter. We have previously shown using that system that human PINK1 S228A fails to phosphorylate Ub Ser65 in cells after inducing mitochondrial depolarization with CCCP (Rasool et al., 2018). Under similar experimental conditions, mutation of the conserved and proximal Lys364 (Lys339 in Tc) and Tyr454 (Tyr429 in Tc) completely abolished the ability of hPINK1 to generate pSer65 Ub chains (Figure 5B). However, testing of distal elements is complicated by poor sequence conservation at the insert 2 interface (αi helix) and part of the A-loop (Figure 5A and Suppl. Figure S6). We thus generated a homology model of the human PINK1 autophosphorylation complex to identify residues potentially critical for dimerization. The model maintains critical interactions proximal to the phosphorylation site, consistent with the involvement of both Lys364 and Tyr454 (Figure 5C). Our model also indicates that Tyr404 would sit in a position similar to Glu379 but would instead form hydrophobic contacts with Cys388 and Leu389 (Figure 5C). The mutation Y404A indeed results in a reduction of Ub phosphorylation in cells (Figure 5B). This is consistent with the observation by the Muqit lab that deletion of insert 2 reduces Ub phosphorylation (Kumar et al., 2017). None of the autophosphorylation interface mutants result in a decrease of PINK1 expression or reduced accumulation in response to CCCP, hence the reduction in pUb is likely the result of the inability of PINK1 to autophosphorylate at Ser228. We conclude that human and insect PINK1 autophosphorylate through a common interface.

**Figure 5.**
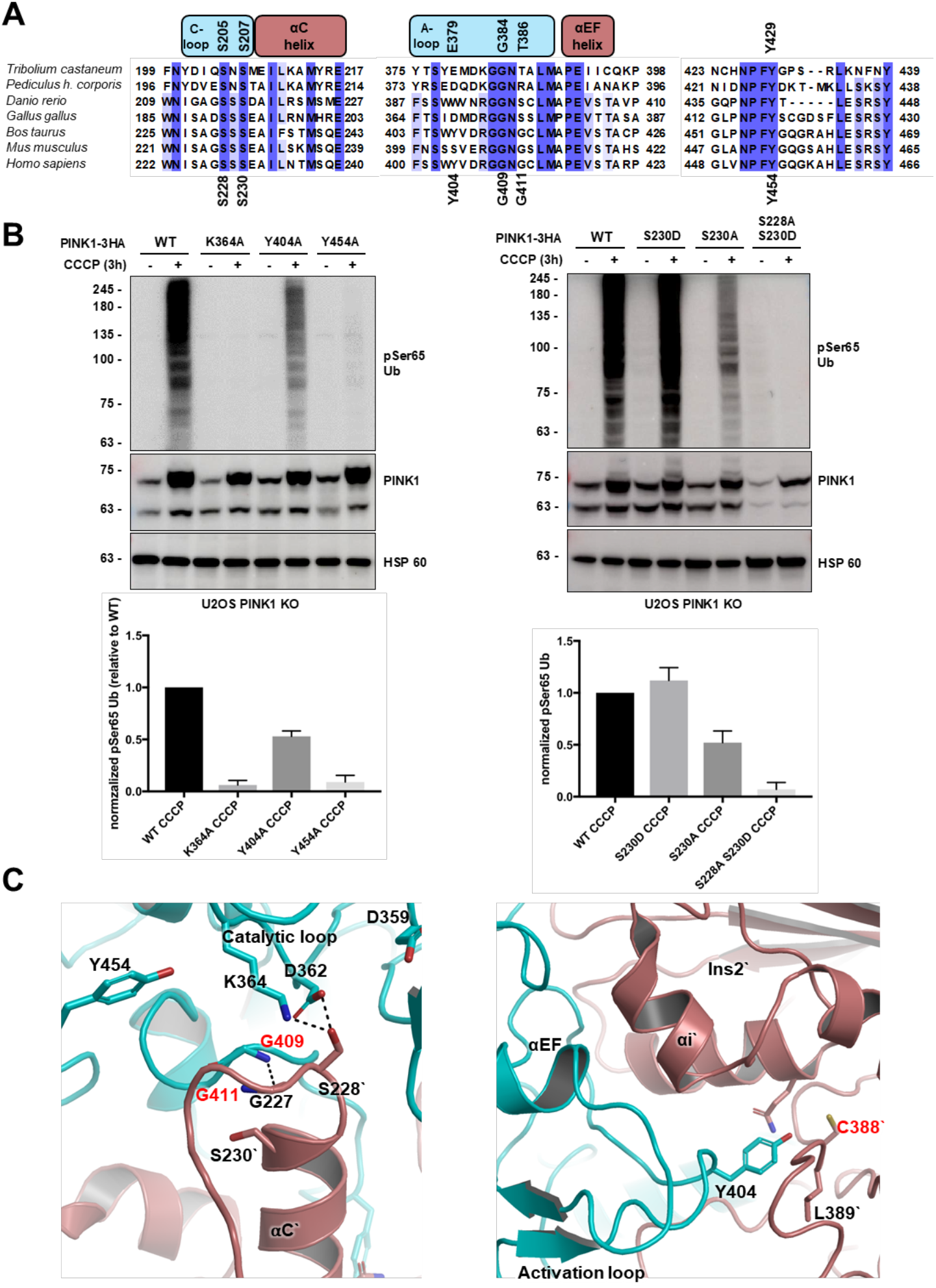
Autophosphorylation is required for activation of Ub kinase activity in human PINK1 in cells. **(A)** Sequence alignment of PINK1 orthologs for the C-loop and αC helix region (left), the A-loop (middle) and the loop connecting the αF and αG regions containing the conserved Y429 (Y454 in human PINK1). **(B)** (top) Immunoblots for pSer65 Ub, PINK1 and HSP60. U2OS PINK1 KO cells were transected with WT-3HA or mutant-3HA (autophosphorylation interface) and treated with 10 µM CCCP for 3 hours. Lysates were analyzed with the indicated primary antibodies. (below) quantification of the level of the relative levels of pSer65 Ub in the CCCP treated samples. The intensities were first normalized relative to the level of PINK1 in the CCCP treated sample and then ratio to level of pSer65 for WT CCCP treated sample was calculated and plotted. Bars represent mean ± SD (n=3). **(C)** The interface at the active site and insert 2 of the human PINK1 autophosphorylation complex homology model. Residues in red correspond to PD mutants.

Human Ser230 is conserved across PINK1 orthologs (Figure 5A). While the equivalent residue is phosphorylated in two of the other insect PINK1 structures (Schubert et al 2017; Kumar et al. 2017), it is unclear whether Ser230 in human PINK1 is also phosphorylated (Okatsu et al. 2012). We therefore mutated human PINK1 Ser230 to either alanine or aspartate to determine the role of this residue in Ub phosphorylation (Figure 5B). The S230D mutation did not significantly alter Ub phosphorylation, in stark contrast with the TcPINK1 S207D mutation which abolished autophosphorylation activity (the S207D mutant could not be phosphorylated at Ser205 in vitro, and thus we could not test Ub/Ubl phosphorylation). This effect could be attributed to differences in the C-loop and A-loop interactions (compare Figure 5C and 2B; see Discussion). On the other hand, the S230A mutation mildly reduced Ub phosphorylation. This data suggest that Ser230 may play a role as a phosphate acceptor, which could replace Ser228 phosphorylation to activate PINK1. However, the double mutant S228A-S230D was completely inactive in Ub phosphorylation, demonstrating that Ser228 phosphorylation remains essential to activate PINK1 in the context of a Ser230 phosphomimetic mutation.

### The NT helix interacts with the C-terminal extension

Our TcPINK1 crystallization construct also included the 32 amino acid NT linker region (a.a. 121-153) linking the kinase domain to the putative transmembrane domain. Deletion of the NT region in TcPINK1 does not impair kinase activity *in vitro* and thus its role remained unclear (Rasool et al. 2018). Our crystal form reveals that the NT linker forms a helix that runs behind the kinase domain and stacks on top of the αK helix in the CTE (Figure 6A). The interface is maintained largely by hydrophobic contacts mediated by Ile133 and Ile137 in the NT helix, and Leu507, Leu508, and Leu515 in the αK helix (Figure 6B). In order to validate that the NT-αK interaction observed in our crystal structure exists in solution and is not an artefact of crystallization, we inserted a 3C protease cleavage site between residues 149 and 150 in our TcPINK1^121-570^ construct and subjected the protein to size-exclusion chromatography (SEC) followed by SDS-PAGE analysis. Cleavage with the 3C protease should not disrupt the interaction between the NT and αK helix and the NT should remain bound to the cleaved kinase domain. Consistent with this hypothesis, we observed a co-migration of the ∼4 kDa NT (a.a. 121-149) with the TcPINK1 a.a. 150-570 fragment produced after cleavage (Figure 6C). We then performed the same analysis with mutants at the NT-αK interface. Consistent with our prediction, the I133E or L508E-T511E double mutant abolished the ability of the NT to co-migrate with the 150-570 fragment and resulted in the NT fragment eluting later (Figure 6C, Suppl. Figure S7). Furthermore, the NT helix has deuterium exchange rates comparable to those observed in the αK helix in the CTE (Figure 6D). The NT helix is therefore an integral part of the C-terminal kinase domain of TcPINK1.

**Figure 6.**
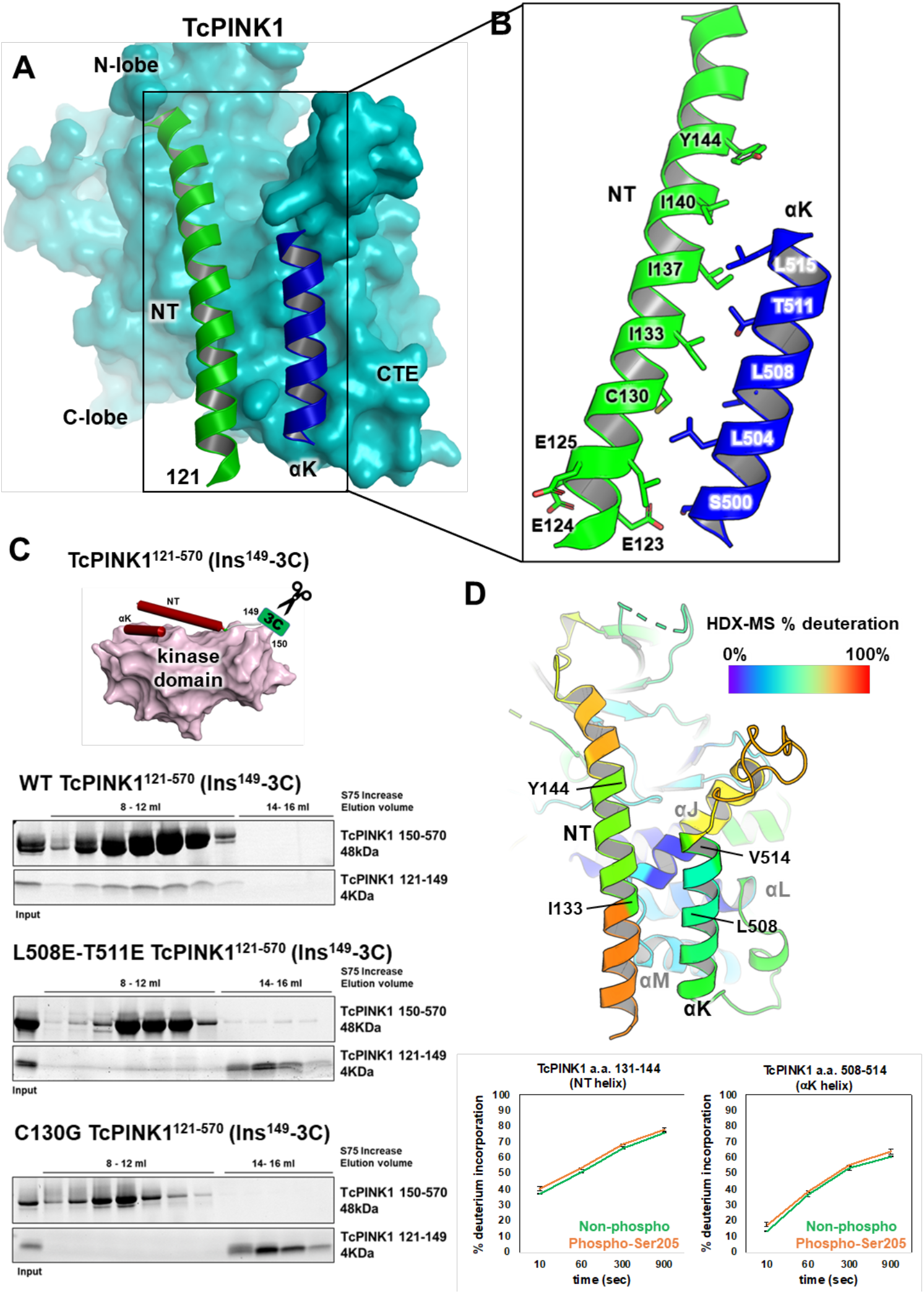
The NT region is helical and interacts with the CTE. **(A)** Structure of the PINK monomer showing the interactions of the NT helix (a.a. 121-149; green) with the αK helix (blue) in the CTE. The remaining regions of the kinase domain are shown as a cyan surface. **(B)** Close-up of residues in the NT and the αK helices. **(C)** NT helix cleavage assays. 3C protease cleavage site was inserted at the end of the NT helix (between residues 149 and 150) for WT and mutants at the NT-αK interface. After 3C cleavage, the protein was subjected to size-exclusion chromatography, and the eluted fractions analyzed by SDS-PAGE and Coomassie staining. The WT shows co-elution of the 4 kDa NT fragment with the TcPINK1^150-570^ fragment. **(D)** % deuterium uptake of peptides from the NT and αK region of the of the kinase for non-phosphorylated and Ser205-phosphorylated WT TcPINK1^121-570^. Bars represent mean ± SD (n=3) (below). The % deuterium uptake was mapped onto the structure of the kinase and color-coded as indicated (above).

The NT helix harbors two PD mutations, C125G and Q126P, which have been reported to reduce the accumulation of PINK1 upon mitochondrial damage and impair Parkin localization to mitochondria (Sekine et al., 2019). Human PINK1 Cys125 is conserved (Suppl. Figure S6) and the equivalent Cys130 in TcPINK1 lies at the interface between the NT and αK (Figure 6B). Gln126 is not conserved and the equivalent Trp131 in TcPINK1 points towards the solvent, but mutation to a proline would disrupt the helical interface regardless. We therefore introduced the C130G and W131P mutations in our 3C-protease TcPINK1 construct to determine their impact on the NT-CTE interaction. Both mutations inhibited the ability of the NT to co-migrate with the kinase domain (Figure 6C, Suppl. Figure S7). This suggests that the PD pathology caused by these mutations results from their inability to maintain the NT-CTE interface.

In human, the N-terminal segment of the NT region contains three glutamate residues (Glu112, 113, 117, or 3E motif) that are important for PINK1 stabilization on mitochondria (Sekine et al., 2019). TcPINK1 also contains three glutamate residues (a.a. 123-125) at the N-terminus of the NT helix (Suppl. Figure S6). In our structure, Glu123 points towards the base of the αK helix, while Glu124-125 are solvent-exposed (Figure 6B). Due to low sequence conservation and differences in length in the NT region between species, it is difficult to establish if these glutamate residues play similar roles. Nonetheless, mutation of these glutamate residues to alanine partially impaired the ability of the NT to bind the kinase domain (Suppl. Figure S7). Thus, one or more of these glutamate residues may help in maintaining the NT helix bound to the CTE.

### The NT-CTE interaction is key for stabilizing PINK1 on mitochondria

In cells, PINK1 is tethered to the TOM complex on mitochondria upon damage, where it dimerizes and autophosphorylates (Lazarou et al., 2012; Okatsu et al., 2013). Given that the 3E motif, which is critical for PINK1 stabilization on the TOM complex, is in the NT helix is in proximity to the CTE, we hypothesized that disrupting the interaction between the NT and αK helices would result in a loss of binding to TOM in response to mitochondrial depolarization. To test this, we selected the A537P variant in the αK helix (Cardona et al., 2011), as well as two highly conserved residues at the NT-αK interface, Ile128 and Leu531 (Figure 7A). Those mutants as well as the autophosphorylation mutants S228A and Y454A were transfected in PINK1 KO U2OS cells and treated with CCCP to induce PINK1 accumulation. The formation of the PINK1-TOM complex was probed using Blue-Native PAGE, as reported previously (Lazarou et al., 2012; Okatsu et al., 2013). WT or S228A PINK1 are able to form this complex, as observed by the formation of a band at 750 kDa (Figure 7B). The autophosphorylation interface mutation Y454A, which also impairs formation of pSer65-Ub, retained the ability to form this complex, showing that autophosphorylation activity is not required for TOM binding. Kinase-dead PINK1 has been also shown previously to form this complex. However, the mutations I128E, L531E, and A537P abolished formation of this complex, implying that the NT-αK interface is critical for TOM complex formation.

**Figure 7.**
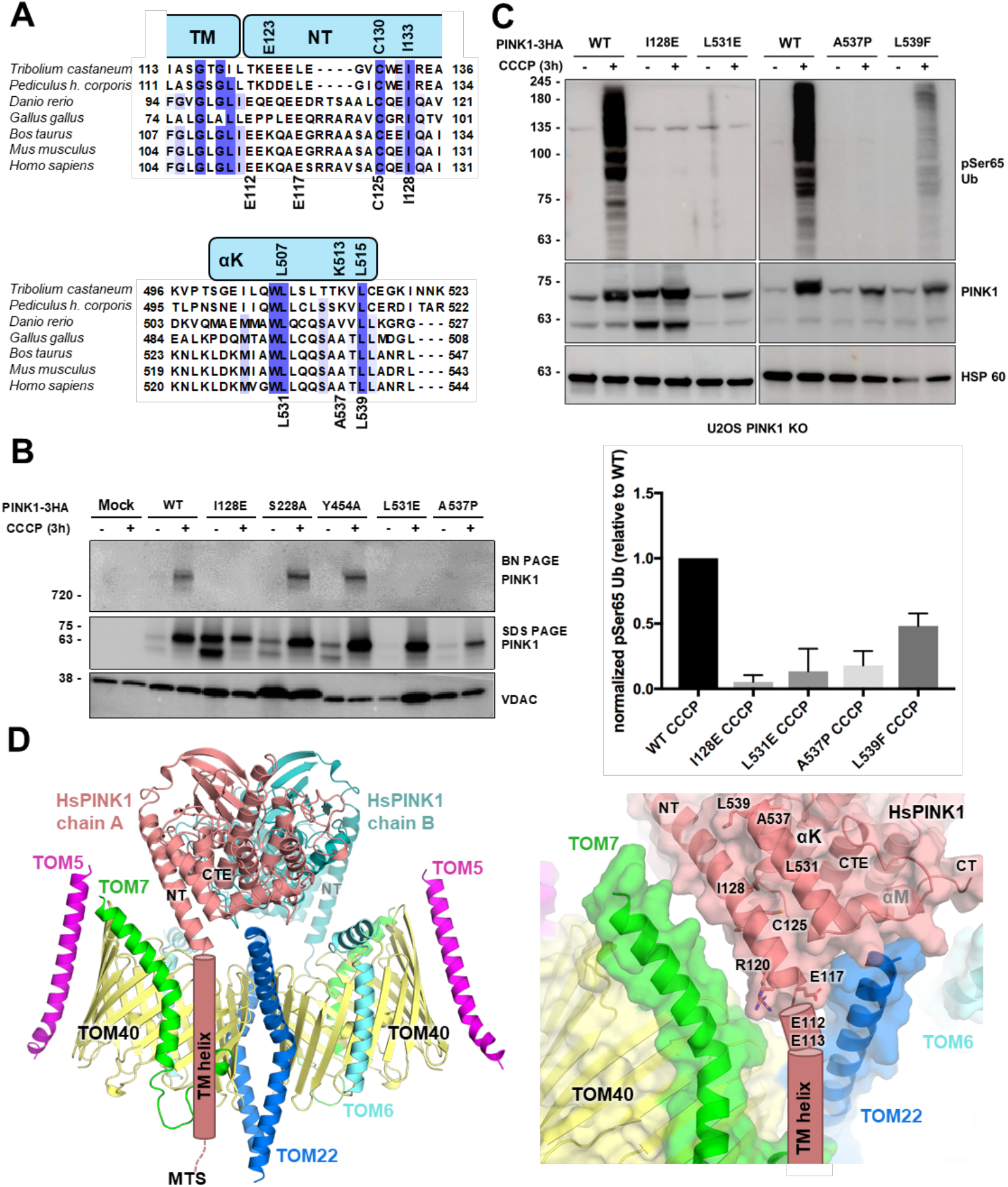
The NT-CTE module and its interaction is essential for stabilizing PINK1 on mitochondria. **(A)** Sequence alignment of the NT (above) and αK helix (below) regions of PINK1. **(B)** Immunoblot of PINK1 (Blue native and SDS PAGE) and VDAC. U2OS PINK1 KO cells were transected with PINK1-3HA WT or mutant and treated with 20 µM CCCP for 3 hours. Mitochondria were extracted, solubilized and analyzed by blotting with the indicated primary antibodies. **(C)** (top) Immunoblots for pSer65 Ub, PINK1 and HSP60. U2OS PINK1 KO cells were transected with WT-3HA or mutant-3HA (NT-αK interface) and treated with 10 µM CCCP for 3 hours. Lysates were analyzed by blotting with the indicated primary antibodies. (below) quantification of the level of the relative levels of pSer65 Ub in the CCCP treated samples. The intensities were first normalized relative to the level of PINK1 in the CCCP treated sample and then ratio to level of pSer65 for WT CCCP treated sample was calculated and plotted. Bars represent mean ± SD (n=3). **(D)** Model of the human PINK1 dimeric autophosphorylation complex docked onto the human dimeric TOM Core Complex (TOM-CC; PDB: 7CK6) (left). (Right) Close-up view of the interface the NT-CTE module and TOM7 and TOM22 subunits of the TOM-CC. The coral cylinder shows the predicted position of E112, E113 and the TM helix of PINK1 (not present in model).

**Figure 8.**
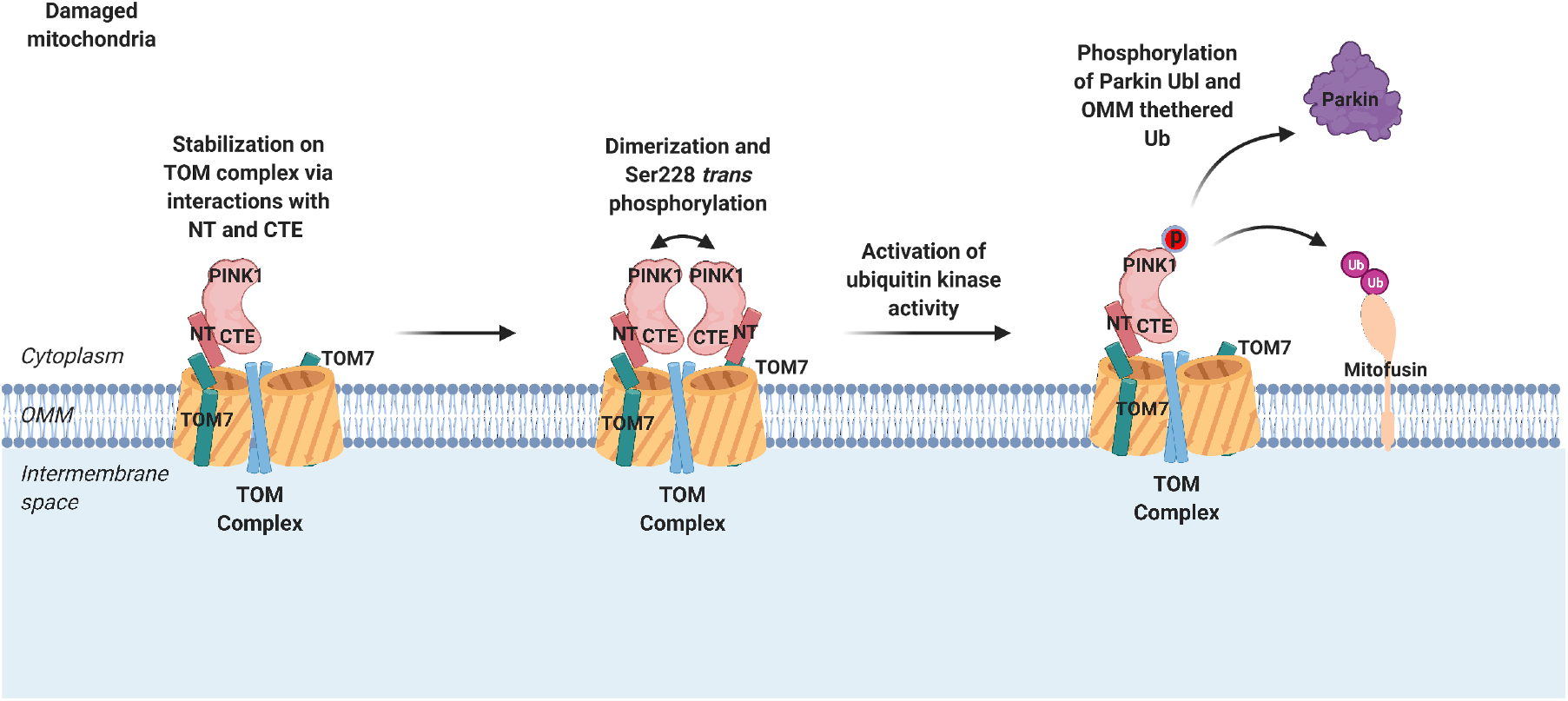
Model of PINK1 stabilization and autophosphorylation on damaged mitochondria leading to Parkin activation. Upon mitochondria damage PINK1 import arrest is stabilized via the interaction of the NT-CTE module of PINK1 with subunits of the TOM complex such as TOM7. Stabilized PINK1 dimerizes for trans-autophosphorylation at Ser228 which is responsible for its ubiquitin kinase activity. PhosphoSer228-PINK1 phosphorylates mitochondrially-tethered Ub and cytosolic Parkin on Ser65 to recruit and activate parkin to ubiquitinate OMM proteins.

To probe the downstream effect of disrupting PINK1-TOM complex formation, we also quantified pSer65 Ub generation by NT-αK interface PINK1 mutants. The I238E, L531E and A537P mutants all inhibited the formation of pSer65 Ub in cells upon mitochondrial depolarization (Figure 7C). Furthermore, the L539F mutant, which was reported in a PD patient (El Haj et al., 2016), partially inhibited pSer65 Ub formation. Human Leu539 is highly conserved and is equivalent to TcPINK1 Leu515 in the αK helix (Figure 6A and Figure 7A). While the αK mutants accumulate in response to CCCP, I128E accumulates in larger amounts compared to WT in the presence or absence of CCCP, but nonetheless fails to produce the 750 kDa complex or generate pUb in response to CCCP (Figure 7 B,C). These results suggest that while these mutants are able to accumulate in response to CCCP, their inability to assemble on the TOM complex renders them unable to autophosphorylate, which reduces phosphorylation of Ub or Parkin.

Since the 3E motif and TOM7 are required for the accumulation of PINK1 on the TOM complex, we posited that the cytosolic fragment of human PINK1 (a.a. 111-581) must interact directly with TOM via the NT helix, in a manner that is conducive to autophosphorylation. To better understand how this might take place, we docked a homology model of the human PINK1 autophosphorylation dimer to the TOM complex. A high-resolution (∼3 Å) cryo-EM structure of the human TOM core complex (CC) was recently reported in both dimeric and trimeric arrangements (Wang et al., 2020). The dimeric structure revealed the arrangements of accessory proteins TOM5, 6 and 7 around the double barrel structure formed by TOM40 and stabilized by TOM22. Since both TOM CC and PINK1 structures have C2 symmetry, we co-aligned their symmetry axes, which allowed us to systematically explore all possible translation and rotation of the PINK1 dimer with its NT helix pointing towards the TOM CC (Suppl. Figure S8A). A first steric search revealed three local minima, but only one led to a stable docking solution after Rosetta refinement (Figure 7D and Suppl. Figure S8B-D). The model reveals the autophosphorylation dimer stretching over of the double-barrel structure, with the NT helix stacking against the N-terminal segment of TOM7. The cytosolic helix of TOM22 is also within interacting distance of the αM helix on the C-terminal end of CTE. This arrangement would position the TM helix in a groove formed by transmembrane segments of TOM7, TOM40 and TOM22, outside the pore. This model is consistent with TOM7’s critical role in stabilizing PINK1 on the TOM complex and would allow PINK1 autophosphorylation at Ser228. Overall, our results show that the NT-CTE interface is critical for formation of the PINK1-TOM complex, which is a required step for PINK1 activation of the Parkin/Ub pathway.

## Discussion

Many kinases become activated through autophosphorylation, either in *cis* or in *trans*. However, PINK1 is unique in its ability to phosphorylate the C-loop in the N-lobe of the kinase domain. For most kinases, autophosphorylation sites typically lie in the A-loop. A survey of published symmetric and asymmetric face-to-face *trans* autophosphorylation kinase dimers (AuroraA, Ire1, PAK1, PAK4) suggests that the αG helix usually mediates critical interactions necessary to facilitate phosphorylation in the A-loop (Suppl. Figure S1C). Interestingly, PINK1 doesn’t have an analogous long αG helix, and the corresponding region of PINK1 is a loop, followed by a short αG helix. In PINK1, it is the αi helix within insert 2 that helps direct phosphorylation to the C-loop. Phosphorylation of PINK1 at the conserved human Ser402 in the A-loop has been suggested based on the inability of the S402A mutation to activate PINK1 and Parkin (Aerts et al., 2015; Okatsu et al., 2012). However, this mutation renders the protein unstable at 37°C, while the S402N mutant, which remains polar but cannot be phosphorylated, functions like WT (Narendra et al., 2013). In our TcPINK1 structure, the equivalent Ser377 is located distal to the active site, and its sidechain forms a hydrogen bond with the backbone amide of Glu379 in the A-loop, which is consistent with this residue playing a structural role rather than a phosphate acceptor role. Thus, our dimer structure provides a firm model to explain why PINK1 uniquely autophosphorylates in *trans* at the C-loop, and not the A-loop.

Ser205/228 phosphorylation is important for the physiological function of PINK1. We and others have shown that human Ser228 is important for substrate phosphorylation in cells following mitochondrial depolarization (Aerts et al., 2015; Kumar et al., 2017; Okatsu et al., 2012; Rasool et al., 2018). Ser228 phosphorylation has also been detected directly by mass spectrometry from PINK1 in human cells (Okatsu et al., 2012). Furthermore, phosphorylation at the equivalent Ser346 in Drosophila has been detected in S2 cells, and PINK1 S346A flies exhibit mitochondrial morphology and motor defects similar to those noted for *PINK1* deletion or kinase-dead strains (Zhuang et al., 2016). By contrast, the S519A mutant strain (human Ser402) showed no defects. This highlights the fundamental role of Ser228 phosphorylation in PINK1 activation, which is conserved across metazoans.

Phosphorylation at human Ser230 has also been suggested to play a role in PINK1 activation (Schubert et al., 2017). However, this residue is not entirely conserved (e.g. Drosophila PINK1 has Ala348 at this site), and we have shown that the human phospho-mimetic S230D mutation cannot restore the activity of the S228A mutant (Figure 5B). It is interesting to note that in the apo structure of TcPINK1, which is phosphorylated at Ser207, Glu205 is retracted and points inwards (Kumar et al., 2017). It is possible that pSer207 or S207D change the conformation of the loop entirely, making Ser205 inaccessible for phosphorylation. However, our dimeric structure and kinase assays show that modification of Ser207 alone disrupts the autophosphorylation interface. Retrospectively, Ser207 phosphorylation, as well as mutations of Thr386 and Ser377 in the AMP-PNP bound structure, might explain why the previously published structures of TcPINK1 did not reveal the autophosphorylation interface. By contrast with TcPINK1, human S230D did not result in a reduction of activity in cells. This could be attributed to differences in the active site and the target serine loop of the two orthologs. Thr386 is indeed replaced by Gly411 in human PINK1 (Figure 5A) and it is possible that the shorter side chain of Gly411 can accommodate an aspartate at the 230 position (Figure 5C). It is interesting to note that G411S has been proposed as a dominant negative PD mutation that impairs the generation of pSer65 Ub (Puschmann et al., 2017). This would be consistent with our dimeric model. However, heterozygous carriers of the mutation do not have a significantly increased incidence of PD, but it cannot be excluded that the mutation has a minor effect on disease risk (Krohn et al., 2020).

In addition to Ser228, Ser230, and Ser402, other phosphorylation sites have been suggested for PINK1. Of particular interest, the Muqit lab detected Thr257 phosphorylation in human PINK1 in cells treated with CCCP (Kondapalli et al., 2012). However, the mutant T257A is still capable of phosphorylating Parkin, and the Morais lab later found that the mutant T257A recruits Parkin to mitochondria (Aerts et al., 2015). Thr257 phosphorylation may therefore simply reflect proxy activation of the kinase. Intriguingly, human Thr257 is equivalent to Glu234 in TcPINK1, which is located in the αi helix (Suppl. Figure S6). In our dimeric human PINK1 homology model, the sidechain of Thr257 points towards the αEF helix in the other subunit (Figure 5C). Phosphorylation at this site would thus modulate dimerization, either by causing a clash which would lead to dissociation of the complex, or by strengthening dimerization through interaction with Arg422 or Arg464. On the other hand, the equivalent Glu231 in the PhPINK1:Ub^TVLN^ structure points away from the substrate (Schubert et al., 2017). Thr257 phosphorylation would therefore not likely affect Ub phosphorylation once Ser228 is phosphorylated, which is the primary activation event.

Our new crystal form is the first to represent PINK1 molecules in a fully primed state for phosphoryl transfer, characterized by the proximity of the target serine to the catalytic residues Asp337 and Asp359, and the closed position of the P-loop (Figure 2B,E and Suppl. Figure S2B). While the modified Ub^TVLN^, which forces slippage of the β5 strand of Ub to expose the Ser65 loop, makes contacts with a.a. 382-385 in the A-loop of the PhPINK1:Ub structure (Schubert et al., 2017), the longer distance from the catalytic aspartate indicates that the Ser65 loop would have to move further into the active site to accept the phosphate (Figure 2B). We predict that the Ser65 containing loop of Ub would make interactions with the catalytic aspartates, as well as Lys339, Tyr429 and the activation loop, in a similar fashion to the Ser205 loop in our structures.

While the non-phosphorylated and phospho-Ser205 TcPINK1 structures we present here have highly similar conformations, the latter likely does not represent the major state in solution. Indeed, the differences observed by HDX-MS between the two forms (Fig. 4D,E and Suppl. Figure S5) suggest more significant conformational changes that prime PINK1 for Ub/Ubl binding. Rather, our phospho-Ser205 structure containing a hydrolyzed AMP-PN molecule in the ATP-binding site (Suppl. Figure S2B,C) represents the ADP-bound state immediately after transfer of the gamma phosphate from ATP to the acceptor Ser205. This complex could serve as a scaffold for the design of small-molecule nucleotide analogs that promote autophosphorylation (Dar and Shokat, 2011; Hertz et al., 2013). Critically, this complex would need to dissociate in order to bind a new ATP molecule, as well as to bind Ub or Parkin since the substrate binding sites overlap (Figure 4A). The precise relationship between the dynamics of the αC helix and the Ser205 loop with nucleotide binding, phosphorylation, insert 3 folding, and substrate binding should be addressed in future structural and biochemical studies.

One of our main findings is the observation that distal elements, in particular insert 2 and its αi helix, are critical for autophosphorylation. This is supported by our mutagenesis experiments (Figure 2D), as well as the inability of PINK1 to phosphorylate a synthetic peptide harboring the C-loop (Suppl. Figure S3). Similarly, PINK1 is unable to phosphorylate a peptide corresponding to the Ser65-containing Ub loop (Lai et al., 2015), which is consistent with the distal Ile44-binding patch being necessary for Ub-Ser65 phosphorylation. We propose that inserts 2 and 3, which are unique features of PINK1 orthologs, have evolved specifically to enable the novel functions of phosphorylation at the C-loop and Ub/Ubl respectively. The role of insert 1, which is divergent between species (Suppl. Figure S6) and is disordered in our structures, remains to be elucidated.

Our study also reveals the structure of the NT helix, its interaction with the CTE, and the role of this module in stabilizing PINK1 on the TOM complex. Interestingly, an NT helix is also found in the Sg269/PEAK1 (Ha and Boggon, 2018) and Sg223/Pragmin (Patel et al., 2017) pseudokinases (referred to as αS or αN1 helix in the literature). These proteins contain a SHED domain (Split Helical Extension Domain), or pseudokinase domain-flanking region, in which the αS helix N-terminal to the pseudokinase domain interacts with the αK helix in a CTE-like domain via hydrophobic contacts. In PEAK1 and Pragmin, the SHED domain has been shown to be essential for creating a back-to-back homodimeric interface in solution even at low protein concentrations. The αS and αK helices run in a criss-cross fashion forming an ‘XX-shaped four-helical bundle’. Our crystallographic contacts also reveal a four-helical bundle between two TcPINK1 subunits. The interface involves the NT and αK helices and has a buried surface area of 2100 Å^2^, which is less than half the area of the autophosphorylation interface. Wild-type TcPINK1^121-570^ is largely monomeric in solution (based on SEC), but we previously observed that TcPINK1^143-570^ migrates as a dimer (Rasool, 2016). Removal of the NT helix would indeed expose hydrophobic residues in the αK helix, which would favor dimerization, as observed in all previous crystal structures that lack the NT helix. While the TcPINK1 back-to-back dimer is likely a crystallization artefact, it suggests that the NT-CTE is a protein-protein interaction module, and we speculate that this reflects PINK1’s ability to make contacts with helical TOM core subunits (5, 6, 7, 22).

Our native PAGE gels and analysis of Ub phosphorylation in cells establish that disruption of the NT-αK interface prevents docking on the TOM complex, which in turn prevents activation of PINK1’s Ub phosphorylation (Figure 7). Since the deletion of the NT helix is dispensable for kinase activity in vitro (Rasool et al., 2018), we posit that mutations in the NT or αK helix do not abrogate human PINK1’s kinase activity per se, but rather interferes with its ability to autophosphorylate at Ser228. Importantly, mutation of the 3E motif in the NT linker, which impairs PINK1-TOM complex formation and Parkin recruitment, still localizes to mitochondria (Sekine et al., 2019). Furthermore, the PD mutants C125G and Q126P also localize to mitochondria, but yet are unable to recruit Parkin (Sekine et al., 2019). These two mutations, as well as I111S, fail to stabilize PINK1 upon mitochondrial depolarization, an effect that could be rescued by silencing of the OMA1 protease, which possibly counteracts TOM7-dependent stabilization (Sekine et al., 2019). Thus, PINK1 activation is depending not only on its mitochondrial localization, but also on the specific positioning of two subunits adjacent to each other within the PINK1-TOM complex.

In our symmetry-driven docking model, TOM7 is juxtaposed to the NT helix, which would position the TM helix outside the TOM40 barrel (Figure 7D). This is consistent with the observation that PINK1 accumulation does not interfere with mitochondrial import through TOM40 (Greene et al., 2012; Lazarou et al., 2012). Since PINK1 is imported through the TOM channel, the implication is that the PINK1 N-terminus must undergo lateral transfer through TOM40. This type of lateral transfer has been shown for outer membrane protein following import through yeast TOM (Harner et al., 2011), as well as through SAM50 for insertion of β-barrel outer membrane proteins (Hohr et al., 2018). Comparison of dimer and trimer assemblies in the cryoEM structures of fungal and human TOM CC show conformational changes that suggest how lateral release could take place through dynamic remodeling of the TOM22 helices and TOM40 β-sheet “unzipping” (Araiso et al., 2019; Bausewein et al., 2017; Wang et al., 2020). The TOM20 and TOM70 proteins are also implicated in binding to PINK1 (Lazarou et al., 2012; Okatsu et al., 2013), but these subunits are missing from the currently available TOM structures. Our dimeric PINK1 structure will facilitate the structural elucidation of the PINK1-TOM complex, which will reveal how TOM20 and TOM70 are positioned with respect to the core complex. In summary, our structure provides a molecular framework for understanding how PINK1 detects mitochondrial damage, a crucial step for the long-term survival of neurons implicated in PD.

## Acknowledgments

We thank colleagues from the Trempe, Gehring, and Fon labs for stimulating discussions and help with the project. The Canadian Macromolecular Crystallography Facility (CMCF) beamlines at the Canadian Light Source (CLS) are supported by the Canada Foundation for Innovation (CFI), the Natural Sciences and Engineering Research Council of Canada (NSERC), the National Research Council of Canada, the Canadian Institutes of Health Research (CIHR), the Government of Saskatchewan, and the University of Saskatchewan. We express our deepest appreciation to the CMCF staff, in particular P. Grochulski, and S. Labiuk. We thank the McGill Pharmacology SPR/MS facility (M. Hancock) and the CFI for support, as well as the Drug Discovery (Nidia Lauzon) and Proteomics (Lorne Taylor, Amy Wong, Jennifer Nedow) platforms at the Research Institute of the McGill University Health Centre (RI-MUHC). This work was supported by a Canada Research Chair (Tier 2) in Structural Pharmacology to J.-F.T., as well as grants from NSERC (#06497-2015), CIHR (#153274), Parkinson Canada (#2017-1277) and the Michael J. Fox Foundation (#12119). S.R. was supported by a studentship from Parkinson Canada, as well as from the Centre de Recherche en Biologie Structurale (Fond de Recherche du Québec - Santé). E.A.F. is supported by a Foundation grant from the CIHR (FDN-154301) and a Canada Research Chair (Tier 1) in Parkinson’s Disease. M.E. is supported by a CIHR Banting Fellowship. S.V. is supported by an FRQS postdoctoral fellowship. G.L. is supported by the CIHR and CFI.

## Author Contributions

Protein purification and crystallization, S.R.; Crystallography and structure determination, S.R., S.V.; Biochemical assays and cell-based assays S.R.; Mitochondrial isolation and BN-PAGE analysis, M.E., E.F.; HDX-MS data collection and analysis, N.S., G.L.; Writing & conceptualization, S.R. and J.-F.T.

## Supplementary figures and legends

**Supplemental Figure S1.**
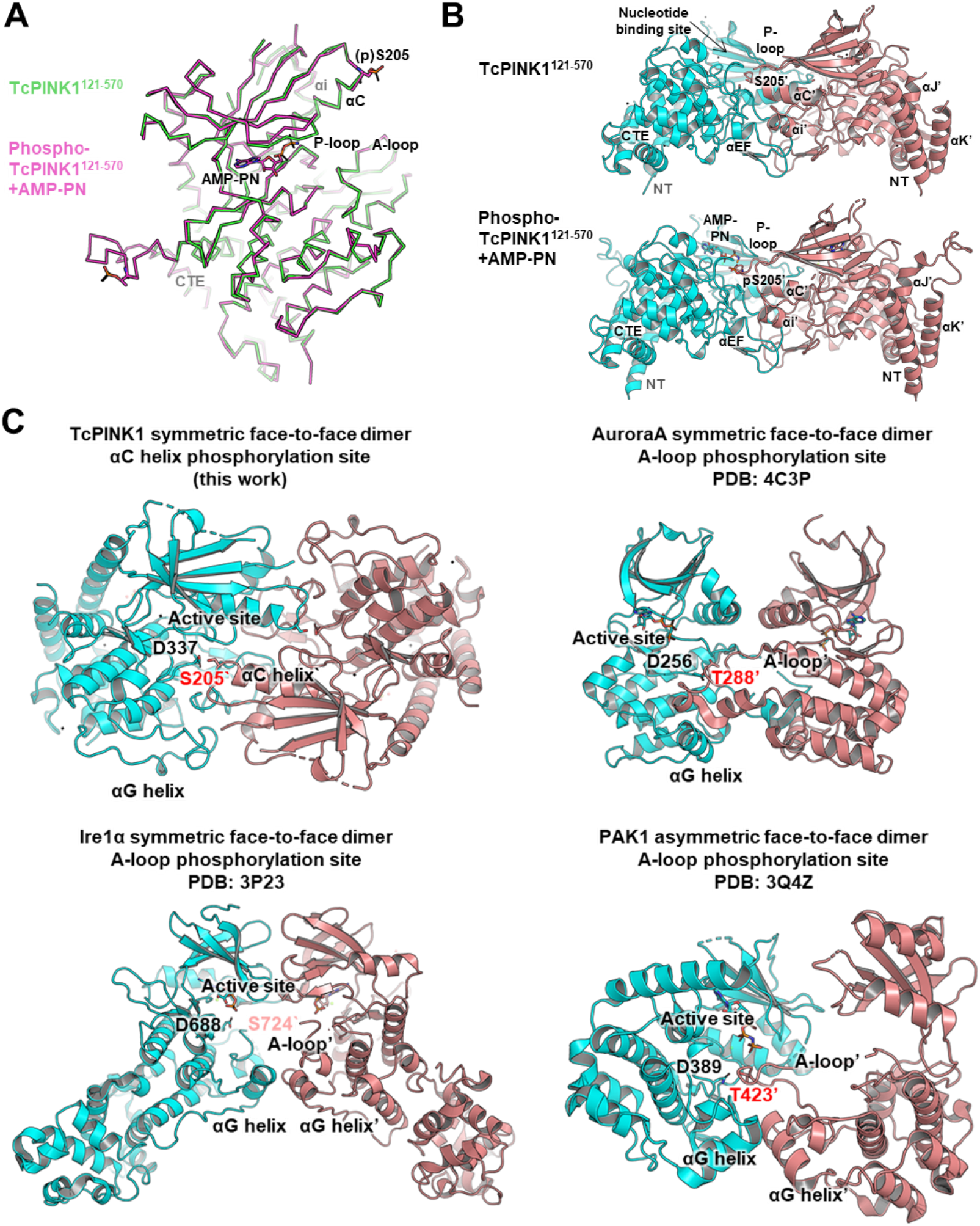
Structure of TcPINK1 autophosphorylation complex, related to Figure 1. **(A)** Ribbon overlay of the AMP-PN-bound Ser205-phosphorylated and the non-phosphorylated monomer structures. The only major difference observed between the structures in the in conformation of the P-loop. **(B)** Side-by-side comparisons of the dimeric autophosphorylation complex of the Ser205-phosphorylated and the non-phosphorylated monomer structure revealed from crystallographic symmetry. **(C)** Comparison of the TcPINK1 dimeric autophosphorylation complex with previously published face-to-face *trans* autophosphorylation complex dimers of other Ser-Thr kinases. PINK1 autophosphorylation complex assembly is primed to target phosphorylation at the C-loop at the tip of the αC helix while for many other kinases autophosphorylation is directed at a conserved serine or threonine in the A-loop.

**Supplemental Figure S2.**
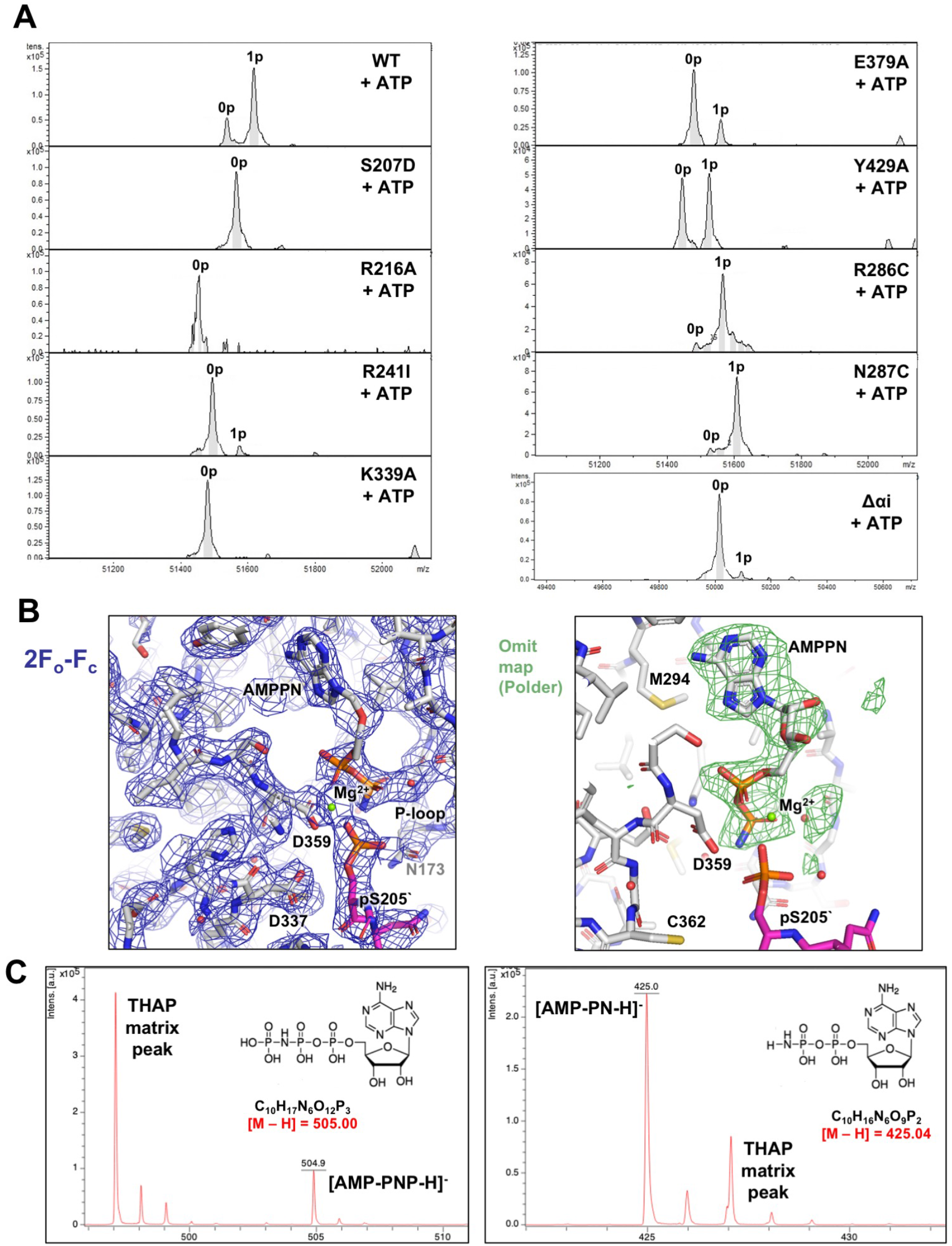
Interactions at the autophosphorylation interface of TcPINK1, related to Figure 2. **(A)** Intact mass spectra of the phosphorylation reactions performed by incubating 1 µM dephosphorylated WT or mutant (autophosphorylation interface) TcPINK1 121-570 with ATP for 30 seconds at 30°C. **(B)** *2Fo-Fc* electron density map (1 σ, left) and Polder omit map (right, 3.5 σ) of the Ser205-phosphorylated structure showing active site residues and the absence of gamma phosphate in AMP-PNP. AMPPN has been seen previously published structure of kinases (PAK4, PDB: 4JDI; PKA, PDB: 4HPT). **(C)** In-vitro hydrolysis of AMP-PNP. MALDI-MS spectra AMP-PNP collected after incubation with TcPINK1 at 288 K for 4 days (duration of crystallization). The sample was spotted on 2’,4’,6’-trihydroxyacetophenone (THAP) matrix.

**Supplemental Figure S3.**
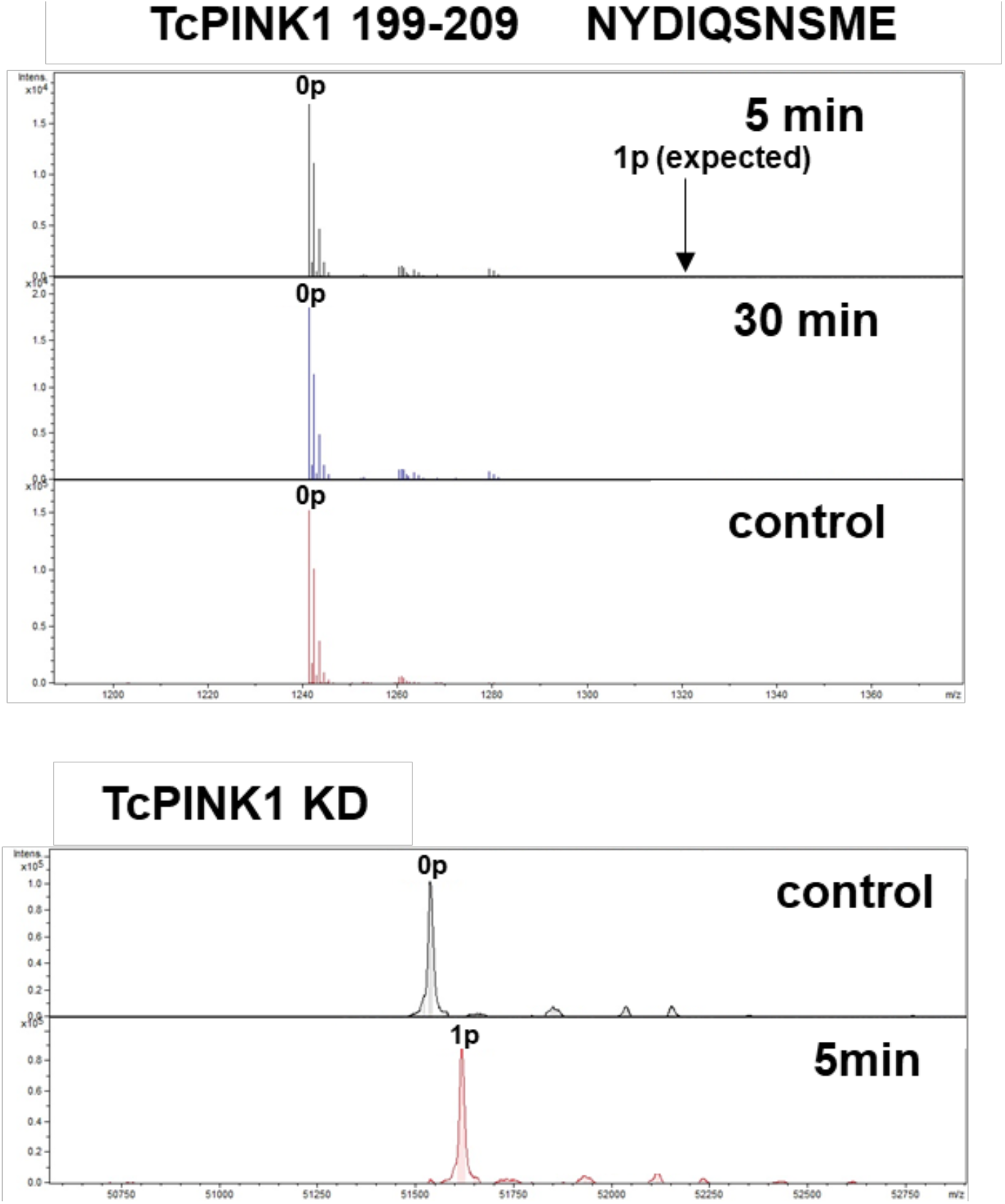
Interactions distal to the active site are essential for trans autophosphorylation, related to Figure 3. Intact mass spectra of the phosphorylation reactions performed using 0.1 µM dephosphorylated GST-WT TcPINK1^121-570^ with 30 µM TcPINK1^199-209^ peptide (containing Ser205) or 30 µM TcPINK1^121-570^ KD for the indicated times.

**Supplemental Figure S4.**
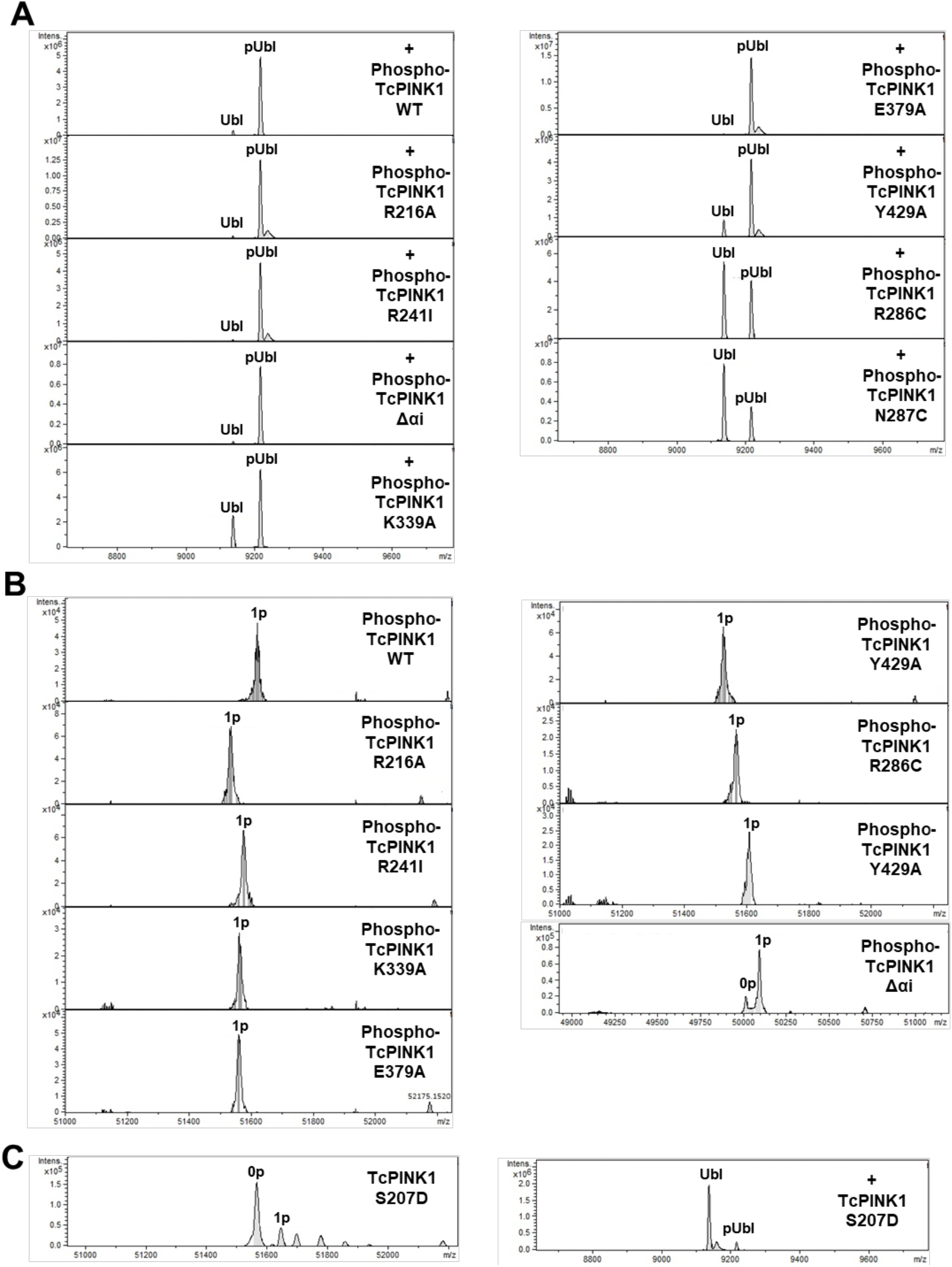
related to Figure 4. Autophosphorylation mutants distal to the active site have no effect on Ubl phosphorylation. **(A)** Intact mass spectra of phosphorylation assays of 60 µM rat Parkin Ubl (1-76) with Ser205-phosphorylated WT or mutant TcPINK1^121-570^. **(B)** Intact mass spectra of TcPINK1 (WT or mutant) used in experiments shown in (A) confirming complete phosphorylation. **(C)** Intact mass spectra of the autophosphorylation and rat Parkin Ubl (1-76) phosphorylation with Ser207D TcPINK1^121-570^.

**Supplemental Figure S5.**
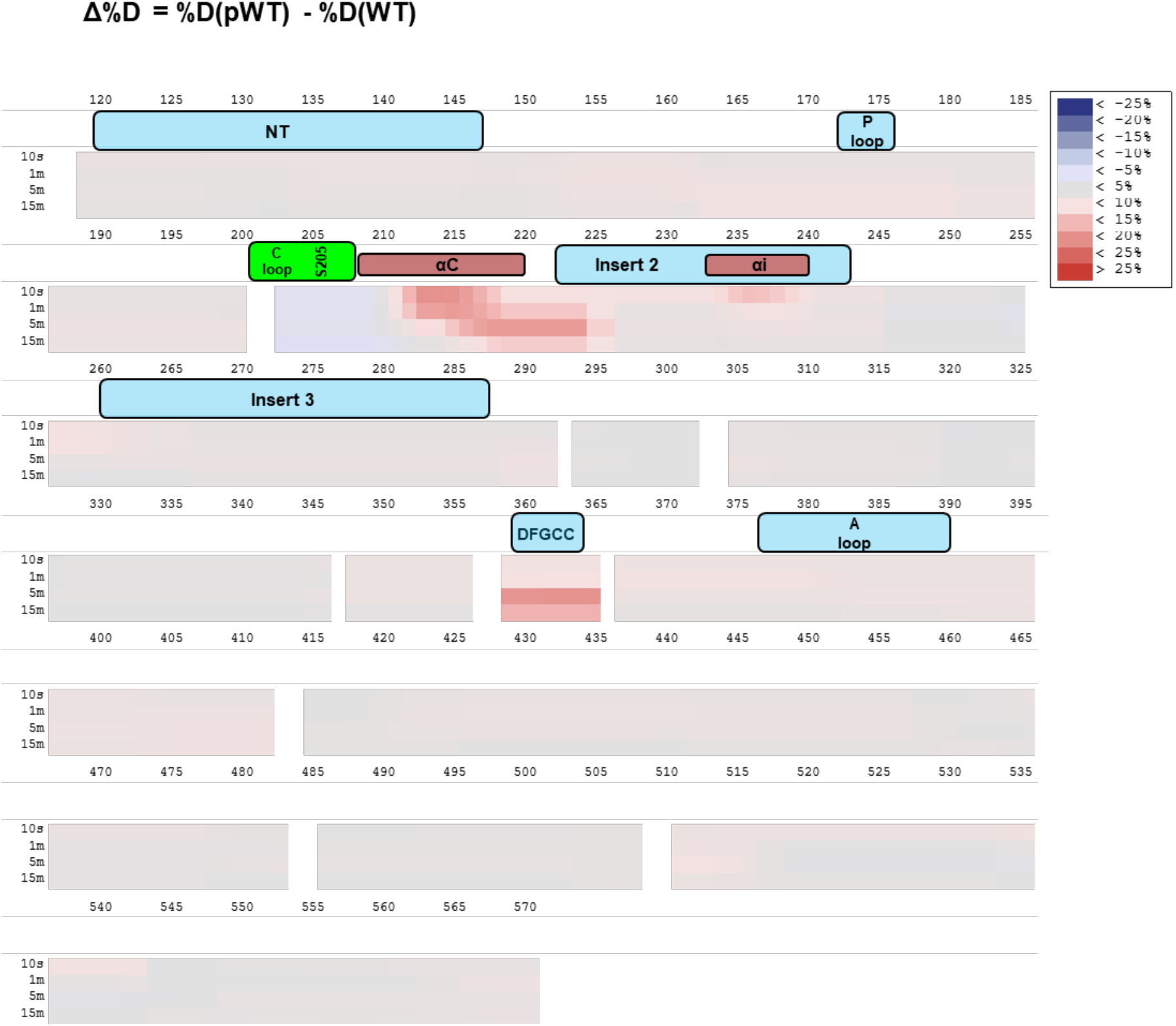
Hydrogen-deuterium exchange mass spectrometry analysis of phosphorylation, related to Figure 4. % change in deuterium uptake between non-phosphorylated and Ser-205 phosphorylated TcPINK1 crystallization construct following different deuteration times, plotted against residue number.

**Supplemental Figure S6.**
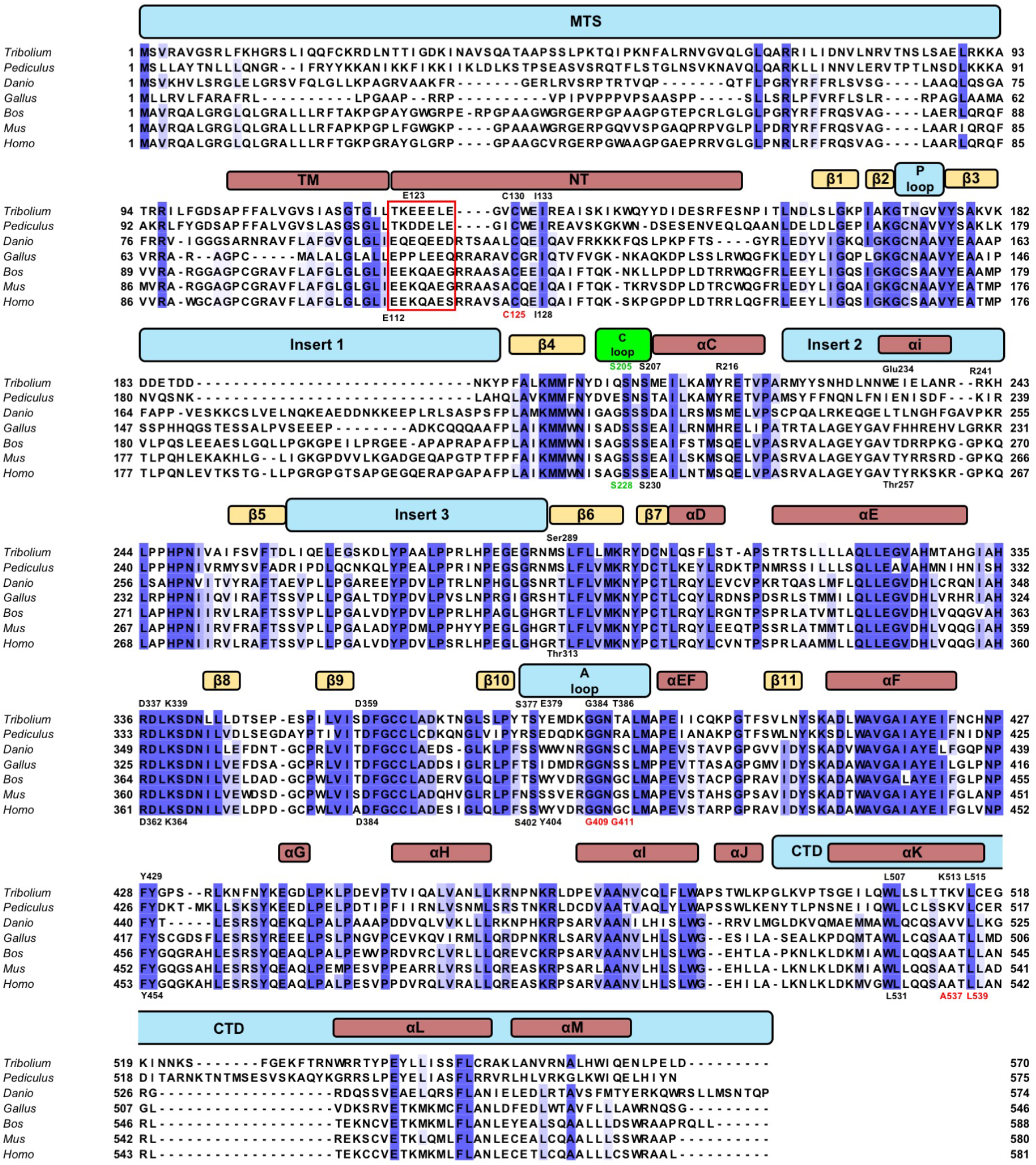
Multiple sequence alignment of PINK1 orthologs, related to Figures 5, 6, 7. Bars in cyan represent different domains or keys regions within the protein, red bars indicate helices while green bars indicate loops. Key residues are indicated. Selected PD mutation sites are colored in red

**Supplemental Figure S7.**
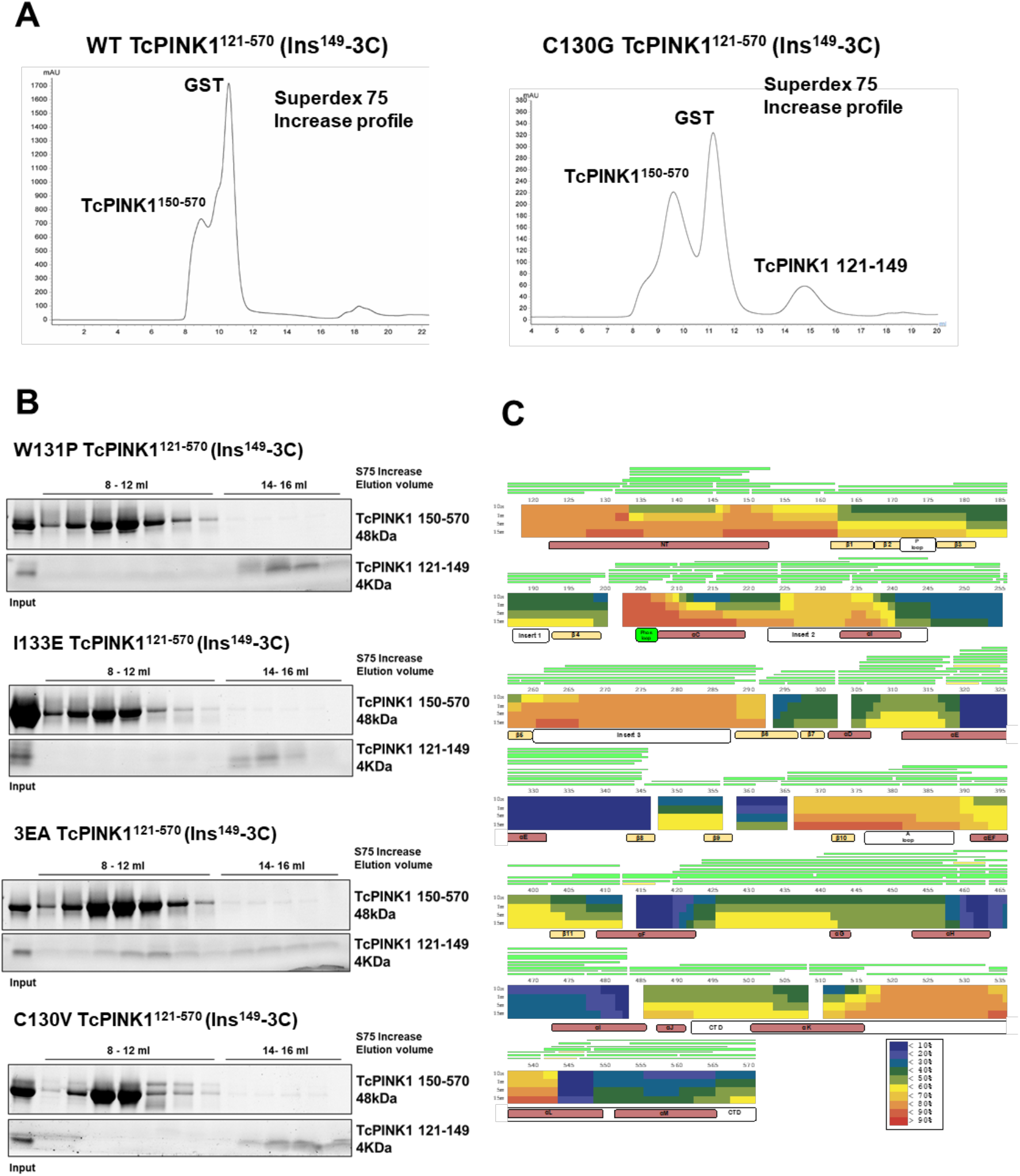
The NT-αK intra-molecular interaction in TcPINK1, related to Figure 6. **(A)** SEC profiles of the cleavage of WT or C130G GST-TcPINK1^121-570^ (Ins^149^-3C). The new peak emerging in 14-16 mL region corresponds to the TcPINK1^121-149^ peptide (confirmed by loading on gel; Figure 6C). **(B)** 3C cleavage reactions of WT or mutant GST-TcPINK1^121-570^ (Ins^149^-3C) loaded on SDS-PAGE after size-exclusion with Superdex75 Increase. **(C)** HDX of TcPINK1^121-570^ at different times after deuteration represented as a heatmap based on the provided color scheme. Key regions of PINK1 are labelled with while bars while helices and sheets are labelled with red and green bars respectively underneath the heatmap.

**Supplemental Figure S8.**
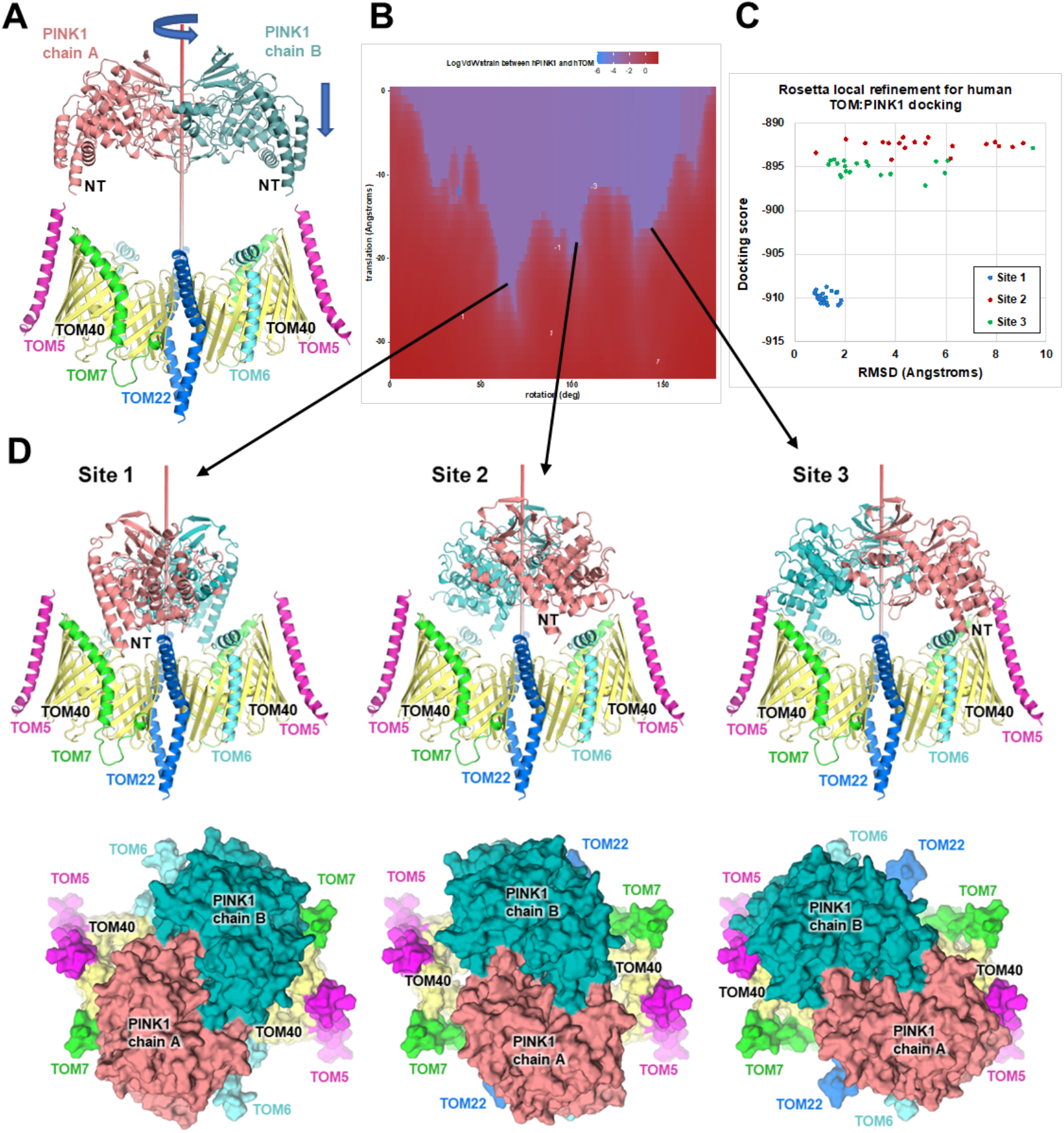
Docking of the human PINK1 homology model to the TOM core complex, related to Figure 7. **(A)** Structures of the human PINK1 dimer homology model (top, this work) and the human TOM core complex (PDB 7CK6). Both structures are aligned with respect to their C2 symmetry axes. **(B)** Plot of the log of the VdW strain as a function of the rotation and translation of the PINK1 dimer. **(C)** Rosetta docking scores and RMSD to the input structures for the 20 lowest energy models generated by the ROSIE docking server, starting from the three sites identified in B. **(D)** Cartoon models of the three site models identified in B, displayed from the side (top) and from the cytosolic side (bottom). Site 1 is the most stable configuration.

**Supplemental Table S1.**
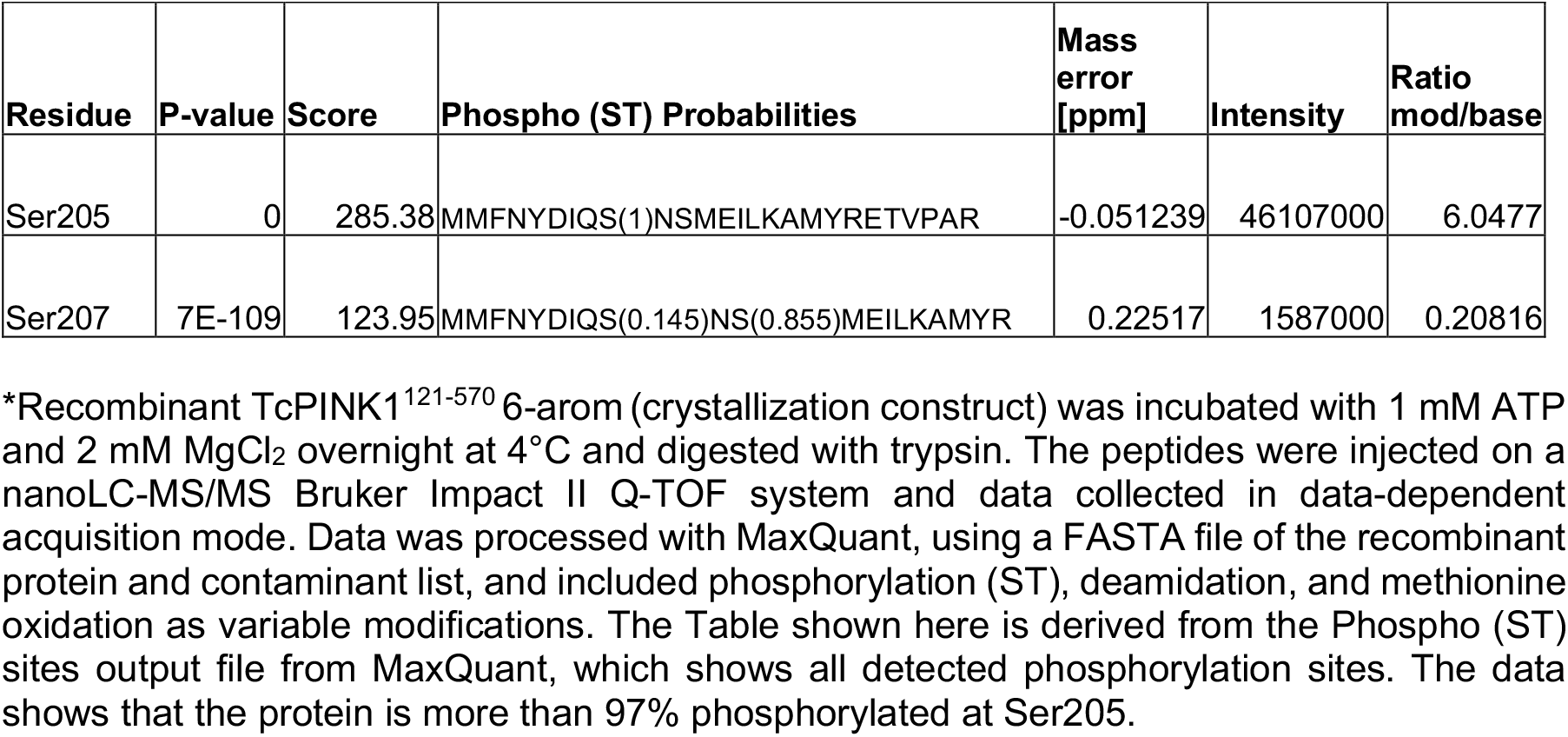
LC-MS/MS statistics for phosphorylated TcPINK1^121-570^, related to Figure 1B.

## METHODS

### RESOURCE AVAILABILITY

#### Lead contact

Further information and requests for resources and reagents should be directed to and will be fulfilled by the Lead Contact, Jean-François Trempe.

#### Materials availability

All plasmids used for expression of insect and human PINK1 are available upon request.

#### Data and code availability

The coordinates and maps for the two structures presented have been deposited in the Protein Databank https://www.rcsb.org: PDB 7MP8, 7MP9.

## EXPERIMENTAL MODEL AND SUBJECT DETAILS

All recombinant proteins were produced in BL21 (DE3) *Escherichia coli* cells. All cloning was performed in DH5-alpha *E. coli* cells. Bacterial cells were propagated in Luria broth (LB) at 37°C with shaking at 180 rpm. Competent cell stocks were prepared in-house for heat shock transformation with plasmids of interests. All experiments with human PINK1 were performed in human bone osteosarcoma epithelial cells (U2OS).

## METHOD DETAILS

### Protein expression and purification

All TcPINK1^121-570^ (WT and mutant) and Parkin Ubl (a.a. 1-76) were expressed as N-terminal GST-tagged proteins from pGEX-6p-1 vector and purified as previously described (Rasool et al., 2018). WT and mutant forms of TcPINK1 used for kinase assays were co-expressed with lambda phosphatase (pET 13S-A co-transformation vector) in BL21 DE3 cells. The Lambda phosphatase plasmid was a gift from John Chodera, Nicholas Levinson, and Markus Seelinger (Addgene #79748). 25 μM MnCl_2_ was added to the cultures after induction with 0.2 mM IPTG to boost the activity lambda phosphatase during protein expression. After elution, the GST was cleaved off using 3C protease overnight at 4°C and the protein was analyzed using intact mass spectrometry to check the phosphorylation state. The proteins were then subjected to size-exclusion chromatography (SEC) in 25 mM HEPES (pH 7.5), 300 mM NaCl, 2 mM MgCl_2_ and 10 mM DTT using a Superdex S200 or Superdex 75 (Cytiva) column in series with GST Trap 4B (Cytiva) to remove GST. The fractions from SEC containing the protein of interest were pooled and concentrated using Amicon-Ultra concentrators (10000 M.W or 3000 M.W. cut-off; EMD Millipore) and frozen at -80 °C until further use.

### Crystallization

Six mutations were introduced in the pGEX-6p-1 TcPINK1^121-570^ plasmid, in non-conserved surface-exposed aromatic residues in different regions of the protein to improve solubility and crystallization. The crystallization construct was co-expressed with lambda phosphatase and purified as described above. To obtain fully phosphorylated protein for crystallization, the eluted protein was cleaved with 3C protease and incubated with 1 mM ATP and 2 mM MgCl_2_ overnight at 4°C, followed by SEC as described above. The phosphorylation state of both non-phosphorylated and phosphorylated protein was confirmed using intact mass spectrometry. Crystals of the non-phosphorylated form were obtained by mixing the 0.3 μl of 2.5 mg/ml protein and 1 mM AMP-PNP with 0.3 μL 0.1 M HEPES (pH 7.0), 20% PEG 4K and 0.15 M ammonium sulfate, at 22 °C using the sitting-drop vapor diffusion method. These crystals were fished and cryo-protected in mother liquor supplemented with 20% PEG400. Crystals of the phosphorylated form were obtained by mixing 0.8 μl of 2.5 mg/ml protein and 1 mM AMP-PNP with 0.8 μL 80 mM MES (pH 6.5), 16% PEG 8K and 0.16 M ammonium sulfate at 15 °C using sitting-drop vapor diffusion method. These crystals were fished and cryo-protected in mother liquor supplemented with 20% PEG400.

### X-ray data collection and structure determination

Diffraction data were collected at the CMCF beamline 08ID-1 at the Canadian Light Source. For the non-phospho TcPINK1^121-570^ structure, 900 images were collected with an oscillation angle of 0.2° at 0.9795 Å wavelength. Reflections were processed with XDSME (Kabsch, 2010) merged, and scaled with *Aimless* (Evans and Murshudov, 2013) For the phospho-TcPINK1^121-570^ structure bound to AMP-PN, 2 datasets of 900 images were collected with an oscillation angle of 0.2° at 0.9795 Å, processed and merged using autoPROC (Vonrhein et al., 2011) and STARANISO (Tickle, 2020). The structures were solved by molecular replacement using the structure of TcPINK1 bound to AMP-PNP (PDB 5YJ9) and refined with the Phenix suite (Liebschner et al., 2019). Models were built with Coot (Emsley et al., 2010). Data collection statistics and refinement statistics are given in Table 1. The non-phosphorylated and phosphorylated AMP-PN-bound structures were deposited to the Protein Data Bank (code 7MP8 and 7MP9, respectively).

### Kinase assays

Autophosphorylation assays of TcPINK1 were performed using 1 µM dephosphorylated WT or mutant protein at 30 °C for 30 sec using 1 mM ATP in 25 mM HEPES (pH 7.5), 300 mM NaCl, 2 mM MgCl_2_ and 10 mM DTT. The reactions were quenched using 0.1% formic acid and immediately transferred to ice and then analyzed using mass spectrometry. For Ubl phosphorylation assays, TcPINK1 or mutants were first completely phosphorylated by incubation with 1 mM ATP up to 10 minutes. K339A displayed weak autophosphorylation activity and had to be incubated overnight at room temperature to obtain complete phosphorylation, while S207D could not be phosphorylated even after incubation with ATP overnight. Phosphorylated TcPINK1 WT or mutant at 1 µM were then incubated with 60 µM Ubl for 5 minutes. The assays were quenched with 0.1% formic acid and immediately transferred to ice and then analyzed using intact mass spectrometry.

### Intact mass spectrometry

Protein solution were diluted in 0.1% formic acid, and 1 µg was injected on a Waters C4 BEH 1.0/10 mm column, washed 5 min with 4% acetonitrile, followed by a 10 min 4-90% gradient of acetonitrile in 0.1% formic acid, with a flow rate of 40 μL/min. The eluate was analyzed on a Bruker Impact II Q-TOF mass spectrometer equipped with an Apollo II ion funnel ESI source. Data was acquired in positive ion profile mode, with a capillary voltage of 4500 V and dry nitrogen heated at 200 °C. Spectra were analyzed using the software DataAnalysis (Bruker). The total ion chromatogram was used to determine where the protein eluted, and spectra were summed over the entire elution peak. Multiply charged ion species were deconvoluted at 10,000-resolution using the maximum entropy method.

### Hydrogen–deuterium exchange mass spectrometry (HDX-MS)

HDX was initiated by diluting 50-150 µM stock solution of TcPINK1^121-570^-6arom or pS205-TcPINK1^121-570^-6arom using 1:15 dilution ratio into the D_2_O-based buffer. The HDX incubation period and temperature were set to 10, 60, 300, 900 sec, and 25°C, respectively. HDX was quenched with chilled quenching buffer (300 mM glycine, 8 M urea in H_2_O, pH 2.4) using 1:4 dilution ratio. Quenched solutions were flash frozen in MeOH containing dry ice, and samples were stored at -80°C until use. For the undeuterated control, initial dilution was made in H_2_O buffer. Prior to ultra-high-performance liquid chromatography (UHPLC)-MS analysis, the deuterated TcPINK1 was digested on an online immobilized pepsin column prepared in-house. Resulting peptides were loaded onto a C18 analytical column (1 mm inner diameter, 50 mm length; Thermo Fisher Scientific) attached to an Agilent 1290 Infinity II UHPLC system. Peptides for each sample were separated using a 5-40% linear gradient of acetonitrile containing 0.1% formic acid for 8 min at a 65 µl/min flow rate. To minimize back-exchange, the columns, solvent delivery lines, injector, and other accessories were placed in an ice bath. The C18 column was directly connected to the electrospray ionization source of the LTQ Orbitrap XL (Thermo Fisher Scientific), and mass spectra of peptides were acquired in positive-ion mode for m/z 200–2,000. Duplicate measurements were performed for each time point. Identification of peptides was carried out in separate experiments by tandem MS (MS/MS) analysis in data-dependent acquisition mode, using collision-induced dissociation. All MS/MS spectra were analyzed using Proteome Discoverer 2.4 (Thermo Fisher Scientific). Peptide searching results were further manually inspected, and only those verifiable were used in HDX analysis. The deuteration (%) as a function of incubation time was determined using HDExaminer 3.2 (Sierra Analytics, Modesto, CA). The first two amino acid residues in peptides were excluded from the analysis.

### NT Cleavage assays

A 3C cleavage site (a.a. LEVLFQGP) was engineered between residues 149 and 150 of TcPINK1^121-570^ WT (pGEX-6p1 plasmid) using PCR mutagenesis and Gibson assembly (NEB). Mutations were then introduced into the NT or αK helix region. The proteins were expressed and purified as described above (without phosphatase co-expression). After elution, the proteins were incubated overnight with 3C protease at 4 °C, resulting in the removal of both GST and the NT. The following day the proteins were resolved by SEC on a Superdex 75 Increase column (Cytiva) equilibrated in 25 mM HEPES (pH 7.5) 150 mM NaCl and 1 mM DTT. The fractions were loaded on a 4-20% PROTEAN stain-free gels and analyzed by fluorescent imaging.

### Homology modelling and docking

Human PINK1 (a.a. 116-581) was modeled using the structure of the non-phosphorylated TcPINK1^121-570^ dimer (this work) as a template with MODELLER, via the PyMod 3.0 suite (Janson and Paiardini, 2020). The symmetry axis of the dimer was aligned with the cartesian Z axis through successive rotations along the X (90°) and Y (60°) axes, based on the orientation of a two-fold symmetry axis in the P6_1_22 space group of the template. The center of mass of the aligned HsPINK1 dimer, determined from a publicly available script (https://pymolwiki.org/index.php/Center_of_mass), was then moved to the origin. The cryoEM structure of the human TOM core complex (PDB 7CK6) was similarly aligned, moved to the origin, and then moved along the Z axis away from PINK1 as a starting point (Suppl. Figure S8A). The Van der Wall (VdW) strain at the interface (VdW_total_ – VdW_PINK1_– VdW_TOM_) was then calculated for every rotation angle (1° increment) and translation (1 Å increment) along the Z axis, using an in-house Python script based on the built-in “*sculpt_iterate*” PyMOL function. The log(VdW) was then plotted in R to identify combination of rotation angles and translation (“sites”) where PINK1 reaches as far down as possible before the VdW strain increases (Suppl. Figure S8B and 8D). Three configurations were selected for refinement in Rosetta using the “docking_local_refine” protocol on the ROSIE server (Lyskov et al., 2013). The docking score and RMSD to the starting model were used to identify a configuration (“site 1”) that gave a stable solution (Suppl. Figure S8C). The lowest energy model from this refinement was used for analysis.

### Cell culture and immunoblotting

U2OS PINK1 KO cell lines were generated by CRISPR in the lab of Edward Fon at the Montreal Neurological Institute. Cells were cultured in DMEM (Dulbeco’s Modified Eagle Medium) supplemented with 10% Fetal bovine serum and 1x Pen/Strep and grown at 37°C. pCMV(d1)TNT PINK1(WT)-3HA was obtained from Noriyuki Matsuda for attenuated PINK1 expression. PCR mutagenesis was used to generate mutations in the kinase domain or the NT of human PINK1. U2OS PINK1 KO cells were transfected with 1 µg pCMV(d1)TNT PINK1(WT)-3HA or mutants in 6-well plates for 24 hours, followed by treatment with 10 µM of CCCP or equal volume of DMSO for 3 hours. Whole cells were harvested, resolved by SDS-PAGE and immunoblotted on the nitrocellulose as described previously (Rasool et al., 2018). The following antibodies were used: PINK1 (Cell Signaling cat# D8G3), HSP60 (Cell Signaling, cat# D307), VDAC (Cell Signaling, cat# 4866S), and phospho-S65 ubiquitin (Millipore EMD, cat# ABS1513). Detection was performed using HRP-conjugated secondary antibody and ClarityTM chemiluminescence (Bio-Rad).

### Mitochondrial isolation and Blue-Native PAGE

For Blue-Native PAGE analysis, U2OS PINK1 KO cells were transfected with pCMV(d1)TNT PINK1(WT)-3HA or mutants. The cells were harvested in mitochondrial isolation buffer containing 20 mM HEPES (pH 7.4), 220 mM mannitol, 70 mM sucrose and protease inhibitor cocktail. Nitrogen cavitation was performed to pellet down mitochondria as described previously (Tang et al., 2017). The mitochondria were then solubilized in a buffer containing 20 mM BIS-TRIS (pH 7.3), 100 mM NaCl, 10% glycerol, protease inhibitors and 1% digitonin and left on a rotor for 3 hours at 4°C. The suspension was spun down at 5000 g for 5 minutes at 4°C and the supernatants were carried forward for Native-PAGE gels. BN-PAGE was performed using Native™ PAGE Running Buffer (Invitrogen) containing 0.002% G-250 (Invitrogen). Gels were shaken in denaturation buffer (10 mm Tris-HCl pH 6.8, 1% SDS, and 0.006% 2-mercaptoethanol) for 60 min after electrophoresis and then transferred to PVDF membranes for immunoblotting. The membranes were blocked with 3% fish skin gelatin (Sigma) in 1 PBS with 0.1% Triton X-100, probed with primary and secondary antibodies, and imaged with an Odyssey infrared imaging system using the manufacturer’s recommended procedures (LI-COR).

## QUANTIFICATION AND STATISTICAL ANALYSIS

Statistics for all crystal structures were determined using software listed above. Statistics generated from data processing and refinement are listed in Table 1. SDS-PAGE gels were processed and analyzed using Fiji (Schindelin et al., 2012). Gel densitometry was performed using a constant box size from which background intensity was subtracted. Graphics for densitometry were generated using Prism 9.

